# Histone Modifications and miRNA Perturbations Contribute to Transcriptional Dysregulation of Hypertrophy in Obstructive Hypertrophic Cardiomyopathy

**DOI:** 10.1101/2024.05.09.593374

**Authors:** Ramin Garmany, Surendra Dasari, J. Martijn Bos, Evelyn T. Kim, David J. Tester, Cristobal dos Remedios, Joseph J. Maleszewski, Keith D. Robertson, Joseph A. Dearani, Steve R. Ommen, John R. Giudicessi, Michael J. Ackerman

## Abstract

**Background:** Recently, we demonstrated transcriptional downregulation of hypertrophy pathways in myectomy tissue derived from patients with obstructive hypertrophic cardiomyopathy (HCM) despite translational activation of hypertrophy pathways. The mechanisms and modifiers of this transcriptional dysregulation in HCM remain unexplored. We hypothesized that miRNA and post-translational modifications of histones contribute to transcriptional dysregulation in HCM.

**Methods:** First, miRNA-sequencing and chromatin immunoprecipitation sequencing (ChIP-seq) were performed on HCM myectomy tissue and control donor hearts to characterize miRNA and differential histone marks across the genome. Next, the differential miRNA and histone marks were integrated with RNA-sequencing (RNA-seq) data. Finally, the effects of miRNA and histones were removed *in silico* to determine their necessity for transcriptional dysregulation of pathways.

**Results:** miRNA-analysis identified 19 differentially expressed miRNA. ChIP-seq analysis identified 2,912 (7%) differential H3K4me3 peaks, 23,339 (21%) differential H3K9ac peaks, 33 (0.05%) differential H3K9me3 peaks, 58,837 (42%) differential H3K27ac peaks, and 853 (3%) differential H3K27me3 peaks. Univariate analysis of concordance between H3K9ac with RNA-seq data showed activation of cardiac hypertrophy signaling, while H3K27me showed downregulation of cardiac hypertrophy signaling. Similarly, miRNAs were predicted to result in downregulation of cardiac hypertrophy signaling. *In silico* knock-out that effects either miRNA or histones attenuated transcriptional downregulation while knocking out both abolished downregulation of hypertrophy pathways completely.

**Conclusion:** Myectomy tissue from patients with obstructive HCM shows transcriptional dysregulation, including transcriptional downregulation of hypertrophy pathways mediated by miRNA and post-translational modifications of histones. Cardiac hypertrophy loci showed activation via changes in H3K9ac and a mix of activation and repression via H3K27ac.

## Introduction

Hypertrophic cardiomyopathy (HCM) is one of the most common genetic heart diseases with significant genetic and phenotypic heterogeneity all contributing to asymmetric left ventricular hypertrophy (LVH) and an increased risk of sudden cardiac death (SCD)^1^. The mechanisms underlying cardiac hypertrophy in HCM remain unclear. Recently, we used a multi-omics approach to characterize the transcriptome and proteome in myectomy tissue from patients with obstructive HCM identifying widespread transcriptional and translational dysregulation^2, 3^.

Interestingly, we observed translational upregulation of several hypertrophy pathways implicated in other forms of cardiac hypertrophy converging on RAS-MAPK signaling, and a counter-regulatory transcriptional downregulation of the same hypertrophy pathways^2^.

Understanding the mechanisms regulating the transcriptional dysregulation in HCM, especially the downregulation of hypertrophy pathways, could uncover new therapeutic targets for HCM. Previous studies in HCM myectomy tissue have demonstrated alterations in miRNA, short non-coding RNA which silence transcript expression^4–9^. Additionally, studies have shown that post-translational modifications of histones, the proteins packaging DNA, contribute to transcriptional dysregulation in heart failure and even activation of hypertrophy pathways in other forms of cardiac hypertrophy^10–17^. In general, increased acetylation of histone H3 subunit lysine residues 9 (H3K9) and 27 (H3K27) enhances gene expression, while hypoacetylation suppresses gene expression^18^. In contrast, trimethylation of the same residues is associated with repression of transcriptional activity (H3K9me3 and H3K27me3, respectively)^19^. Finally, trimethylation of lysine 4 on H3 (H3K4me3) marks active promoters^19^.

Using a combined multi-omics approach in one of the largest collections of HCM myectomy tissue, we sought to characterize the role of miRNA and these five histone modifications on the HCM transcriptome and determine whether these mechanisms may regulate or modify cardiac hypertrophy pathways in HCM.

## Methods

### Cohort Selection and Sample Acquisition

As previously described^2^, patients from this Mayo Clinic IRB-approved (#811-98) study underwent surgical myectomies for symptomatic, obstructive HCM (N = 97). Myectomy tissues were flash frozen immediately and stored at -80C following resection. Control samples (n = 23) were obtained from cardiac donors for which a suitable recipient was not identified. RNA-sequencing and miRNA-sequencing were performed on all samples, while chromatin immunoprecipitation sequencing (ChIP seq) was performed on a subset of the HCM samples (n = 40) and all the control samples (n = 23). Basic demographic information is summarized in **Table 1**.

**Table 1.** Basic Demographic Features of Cohort.

| Cohort Demographics |  |  |  |  |
| --- | --- | --- | --- | --- |
|  | **HCM Total Cohort**<br>**(n = 97)** | **HCM ChIP-seq Cohort**<br>**(n = 40)** | **Control Cohort**<br>**(n = 23)** | **p-value\*** |
| **Female, n (%)** | 44 (45) | 18 (45) | 10 (43) | 0.9 |
| **Average Age at Acquisition, years** | 47 ± 18 | 48 ± 18 | 43 ± 12 | 0.5 |
| **Average Age at Diagnosis, years** | 41 ± 20 | 42 ± 19 | NA | 0.7 |
| **Family Hx of HCM, n (%)** | 34 (35) | 10 (25) | NA | 0.3 |
| **Family History SCA, n (%)** | 14 (14) | 3 (8) | NA | 0.3 |
| **SCA, n (%)** | 1 (1) | 0 | NA | 0.9 |
| **ICD, n (%)** | 27 (28) | 11 (28) | NA | 0.9 |
| **LVWT Max at Diastole, mm** | 23 ± 6 | 23 ± 8 | NA | 0.9 |
| **LV Mass Index, g/m<sup>2</sup>** | 189 ± 79 | 184 ± 89 | NA | 0.7 |
| **Max LVOT gradient, mmHg** | 86 ± 39 | 84 ± 36 | NA | 0.8 |

HCM, hypertrophic cardiomyopathy; Hx, history; ICD, implantable cardioverter defibrillator;
LV, left ventricle; LVOT, left ventricle outflow tract obstruction; LVWT, left ventricular wall
thickness; SCA, sudden cardiac arrest. \*Comparison between HCM total cohort and HCM ChIP-seq cohort.

### RNA-sequencing and miRNA-sequencing

RNA was extracted as previously described using the RNeasy Plus Universal Mini Kit (Qiagen) with modifications per RNeasy Mini Handbook to allow extraction of total RNA, including miRNA^2^. TruSeq Stranded mRNA Prep (Illumina, San Diego, CA) was used for library generation for RNA-sequencing and TruSeq Small RNA Prep (Illumina, San Diego, CA) was used to generate libraries for miRNA. RNA sequencing (RNA-Seq) was performed on Illumina Nov Seq S2 with randomization of batches to reduce confounding. Samples were aligned to GRCh38 using STAR Alignment. Ensembl release 78 was used for gene annotations. Following alignment, secondary analyses, and quality control, the edgeR package (Version 3.28.1) was used for obtaining differentially expressed genes (DEGs) between HCM and controls. mRNA data is deposited under GSE249925 and miRNA is deposited under GSE249757 in the GEO database.

### Differential expression analysis

Gene expression differential expression analysis was re-performed on our previously published^2^ mRNA-sequencing data set using EdgeR software (version 3.32.1). In brief, sample wise raw read counts of genes were normalized using trimmed means of M-values method. Normalized read count data was filtered to remove genes that do not have at least 50 counts in half of the samples of at least one cohort. Normalized counts for the remaining genes were compared between any two experimental groups using generalized linear models configured with negative binomial distribution. Differential expression p-values were corrected using Benjamini-Hochberg method. Genes with an adjusted differential expression p-value of ≤ 0.05 and an absolute log2 fold change of ≥ 1 (where 0.0 represents no change) were considered as statistically significant for further analysis.

The miRNA data was processed using the same edgeR software described above to derive differentially expressed miRNA between any two groups. In brief, sample wise miRNA read counts were normalized as described above. Next, miRNA data were filtered to remove any miRNA that do not have at least 20 read counts in at least half of the samples of single group.

Normalized read counts of remaining miRNA were compared between any two groups and resulting differential expression p-values were corrected using the same method described above. The miRNA with differential expression Benjamini-Hochberg p-value of ≤ 0.05 and an absolute log2 fold change of ≥ 1 (where 0.0 represents no change) were considered as statistically significant for further analysis.

### Principal Component Analysis (PCA)

PCA analysis was performed in R using normalized read counts (version 4.1.3).

### miRNA integration with mRNA data

Two different methods (a supervised method and an unsupervised method) were used to integrate miRNA with mRNA data obtained by comparing any two groups. For supervised integration, statistically significant, differentially expressed mRNA and miRNA were uploaded into Ingenuity Pathway Analysis software (Qiagen). IPA was instructed to use experimentally verified mRNA-miRNA interactions and the directionality of miRNA and mRNA fold changes observed in the group comparison to associate candidate miRNAs with mRNAs. Resulting data were subjected to canonical pathway analysis in IPA and pathways with a -log (Benjamini-Hochberg p-value) ≥ 1.3 were considered for interpretation.

For unsupervised analysis, normalized read counts of statistically significant, differentially expressed miRNA (log2fc ≥ |0.5|) and all transcripts identified in cardiac tissue were loaded into R programming environment (version 4.0.1). Each miRNA was correlated with each mRNA using Spearman method and resulting p-value was corrected using Benjamini-Hochberg method. All miRNA-mRNA pairs with an adjusted correlation p-value of ≤ 0.05 and have a correlation value of ≤ -0.5 (i.e., anticorrelating) were loaded into Cytoscape software (version 3.9.1).

### Chromatin immunoprecipitation sequencing (ChIP-seq)

The Epigenomics Development Laboratory (Mayo Clinic) performed all the procedures from initial chromatin preparation up to quality control measurement and library preparation. About 10 mg – 100 mg of human heart tissues was used for the assay. The tissue was homogenized in 1X PBS, and homogenized tissue was cross-linked to final 1% formaldehyde, quenched with 125 mM glycine, and washed with 1X TBS. The fixed tissue was resuspended with cell lysis buffer (10mM Tris HCl, pH7.5, 10 mM NaCl, 0.5% IGEPAL), incubated on ice for 5 min, homogenized, and centrifuged at 21,130 xg for 2 min. After removal of supernatant, the lysate was washed with MNase digestion buffer (20 mM Tris-HCl, pH7.5, 15 mM NaCl, 60 mM KCl, 1 mM CaCl2) and incubated in the fresh 250 μL MNase digestion buffer in the presence of 500 gel units of MNase (NEB, Cat.# M0247S) at 37°C for 20 min with continuous mixing in thermal mixer. After adding 250 µL of 2X Stop/ChIP/Sonication buffer (100 mM Tris–HCl, pH8.1, 20 mM EDTA, 200 mM NaCl, 2% Triton X-100, 0.2% Sodium deoxycholate), the lysates were sonicated for 15 min (30 s on / 30 s off) using a Diagenode Bioruptor pico and centrifuged at 21,130 xg for 10 min. The supernatant was collected to a new tube, and the pellet was resuspended with 500 μL of 1X ChIP buffer (including final 0.1% SDS), sonicated for 15 min (30 s on / 30 s off) using Diagenode Bioruptor pico, and centrifuged at 21,130 xg for 10 min. The supernatant was collected, and the combined supernatants were utilized as chromatin input.

Subsequent ChIP and library preparation were performed as previously described^20^. The following antibodies were used in the experiment: Anti-H3K4me3 (Diagenode cat.# C1541003); H3K9ac (EDL, lot 1); H3K9me3 (Diagenode cat.# C15410056); H3K27ac (CST cat.# 8173); H3K27me3 (CST, cat.# 9733). The ChIP-seq libraries were sequenced to 51 base pairs from both ends on an Illumina Next-seq or Nova-seq platform in the Mayo Clinic Center for Individualized Medicine Medical Genomics Facility.

### Processing of ChIP-sequencing Data

First the reads underwent adapter trimming using cutadapt^21^ (v0.4.4) followed by aligning to human reference genome GRCh38 (hg38) using Burrows-Wheeler Aligner^22^ (BWA, v0.7.17). Reads pairs were retained as long as one end uniquely mapped with quality of 30 or higher for peak calling. Duplicate reads were removed using Picard (v1.67). Model-based analysis of ChIP-seq (MACS2) package^23^ (v2.0.10) and spatial clustering approach for the identification of ChIP-enriched regions (SICER) package^24^ (v0.1.1) were used to identify narrow and broad peaks respectively using a threshold of FDR ≤ 0.01 and a fold-change of ≥ |2|. Peaks corresponding to blacklisted areas were removed^25^. ChIP-seq signal intensity files were generated in bedGraph format using MACS2 and then converted to the BigWig format using the wigToBigWig command. Finally, differential peaks were identified using diffBind with a default threshold of FDR ≤ 0.05^26^. Differential binding peaks were annotated and pathway analysis performed using ChIPseeker^27^.

### Integrating ChIP-seq and RNA-seq Data

An inhouse script written in R programming language (version 4.0.1) was utilized to integrate the differentially abundant chromatin binding peaks with differentially expressed mRNA data. First, differential peak data from all enrichment experiments (i.e., H3K27ac, H3K27me3, H3K4me3, H3K9ac, and H3K9me3) was loaded and each mark was assigned a sign (+1 or -1 based on whether the mark is an enhancer or repressor). Next, peak marks were annotated and all peaks that are between -1500bp<=TSS<=300bp (i.e., promoter) of any gene were selected. In parallel, RNA differential expression data was loaded. Next, differential RNA expression data and peak abundance were merged based on whether the peaks were within the promoters of the genes (as described above). This yielded three different lists: a) Peaks that are within promoters of expressed mRNA b) Peaks that are within the promoters of non-expressed mRNA and c) mRNA with no peaks in their promoters. Each of these lists were processed separately to understand the contribution of differential chromatin binding to mRNA expression. Each peak-mRNA pair in the first list was further processed by computing a rank score that took into account p-value and fold change of the peak and mRNA as well as its repressor/enhancer status (computed as -1*log2(mRNA differential p-value)*sign(mRNA fold change)+[-1*log2(peak differential p-value)*sign(peak fold change)*(repressor/enhancer status)]). The rank score had a range above zero if the peak-mRNA pair was concordant (i.e., mRNA change was in concordant with the function and directionality of the bound peak) and it had a range below zero otherwise. The rank score also represented the degree of concordance or discordance, with higher positive values representing more concordance whereas lower negative values representing more discordance. The resulting peak-mRNA data was separated into concordant and discordant changes. The integrated concordant data were filtered using appropriate adjusted differential p-value (i.e., using peak’s adjusted differential abundance p-value for integrated peak-mRNA pairs and lone peaks with no mRNA whereas RNA’s adjusted differential expression p-value for genes with no peak in promoters) and further analyzed using Ingenuity Pathway analyses.

## Results

### Differentially expressed miRNA in HCM regulate inflammatory and hypertrophy pathways

In total, 479 miRNAs were detected in the myectomy tissue and as shown in PCA analysis, there was a distinct “miRNA signature” that could almost completely distinguish HCM (N = 97) from control samples (N = 23; **Figure 1A**). Nineteen (4%) miRNAs were differentially expressed between HCM and control samples (log2fc ≥ |1| and FDR ≤ 0.05; **Figure 1B and supplemental excel file**). The miRNA target filtering function of Ingenuity Pathway Analysis was used to integrate the differentially expressed genes (DEGs) and identify the mRNA impacted by these 19 miRNAs **(Figure 1C**). In total, the 19 miRNAs were associated with the expression of 365 DEGs (**Figure 1D**). Pathway analysis of the 365 genes identified 134 pathways impacted by these 19 differentially expressed miRNA, including downregulation of fibrotic, inflammatory (**Figure 1E**) and hypertrophy pathways (**Figure 1F**). Of note, 7 miRNAs (miR-139-5p, miR-100-5p, miR34c-5p, miR-143-5p, miR-21-5p, miR-3656, and miR-148a-3p) were implicated in regulation of the hypertrophy pathways (**Supplemental Figure 1A**).

**Figure 1.**
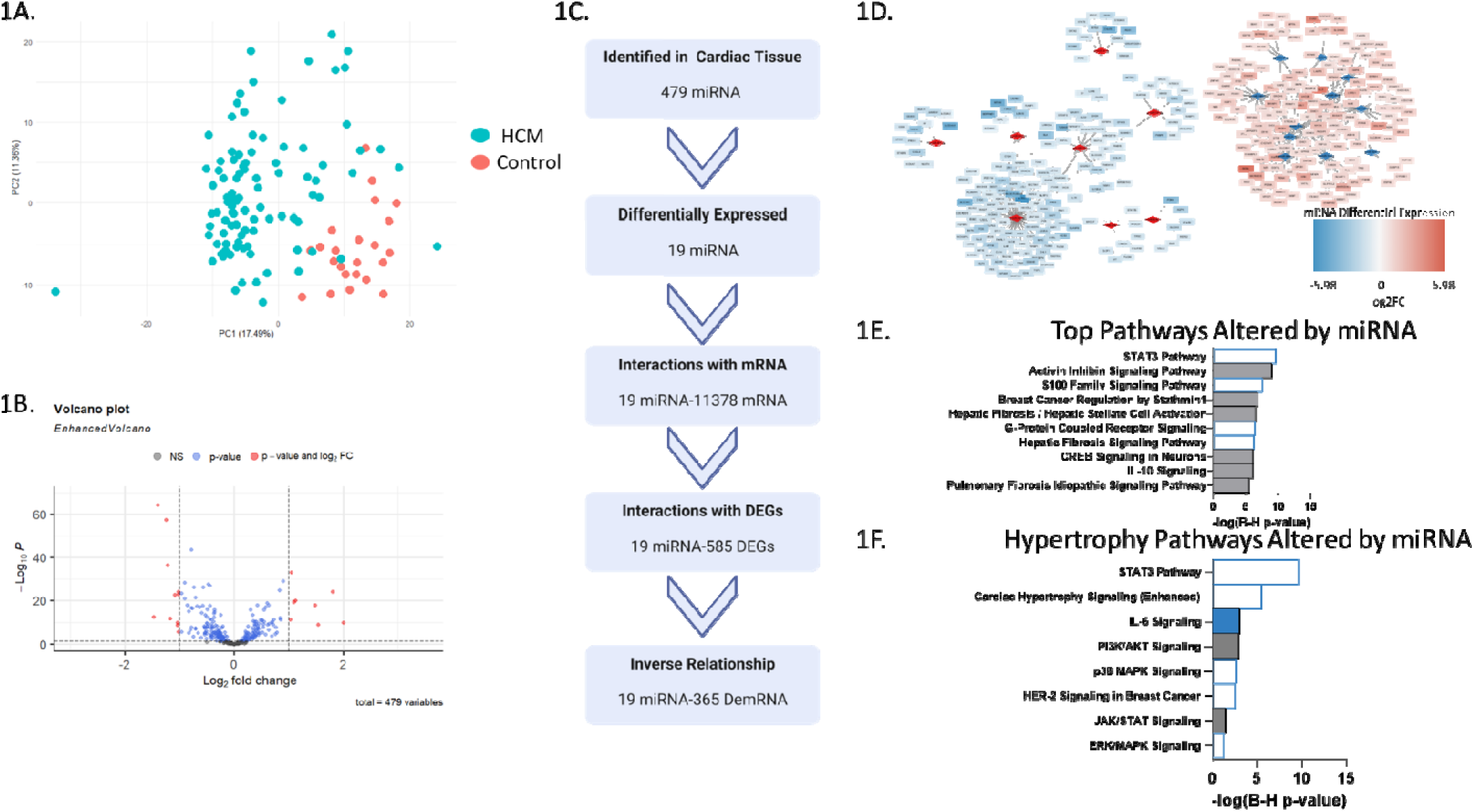
miRNA integration with transcriptome using Ingenuity Pathway Analysis. A) PCA plot comparing miRNA between HCM and controls. B) Volcano plot of miRNA differential expression analysis. C) Workflow to integrate miRNA-sequencing data with RNA transcriptome using Ingenuity Pathway Analysis Method. D) mRNA-miRNA association network using Ingenuity Pathway Analysis integration. E) Pathways most statistically altered by miRNA. F) Hypertrophy pathways regulated by miRNA. Log2FC, log2 fold change.

To validate these findings which rely on previously established mRNA-miRNA interactions, an unbiased correlation approach was also utilized to associate the miRNA with the transcriptome (**Figure 2A**). First, a less stringent cutoff of log2fc ≥ |0.5| was used to identify 79 miRNA that are slightly altered. Next, correlation analysis was performed assessing the association between the 79 miRNAs with all mRNAs across the HCM samples to identify significant miRNA-mRNA pairs (correlation coefficient of ≤ -0.5 and FDR ≤ 0.5). This identified 603 miRNA-mRNA interactions comprised of 40 miRNA and 396 mRNA generating a novel HCM specific miRNA-mRNA associative regulatory network (**Figure 2B**). Pathway analysis revealed the miRNA were most strongly associated with regulation of cardiac hypertrophy and fibrotic pathways (**Figure 2C**) and these genes played a role in important cardiac functions such as positive regulation of heart contraction, heart rate, cardiac muscle contraction, and cardiac conduction (**Figure 2D).**

**Figure 2.**
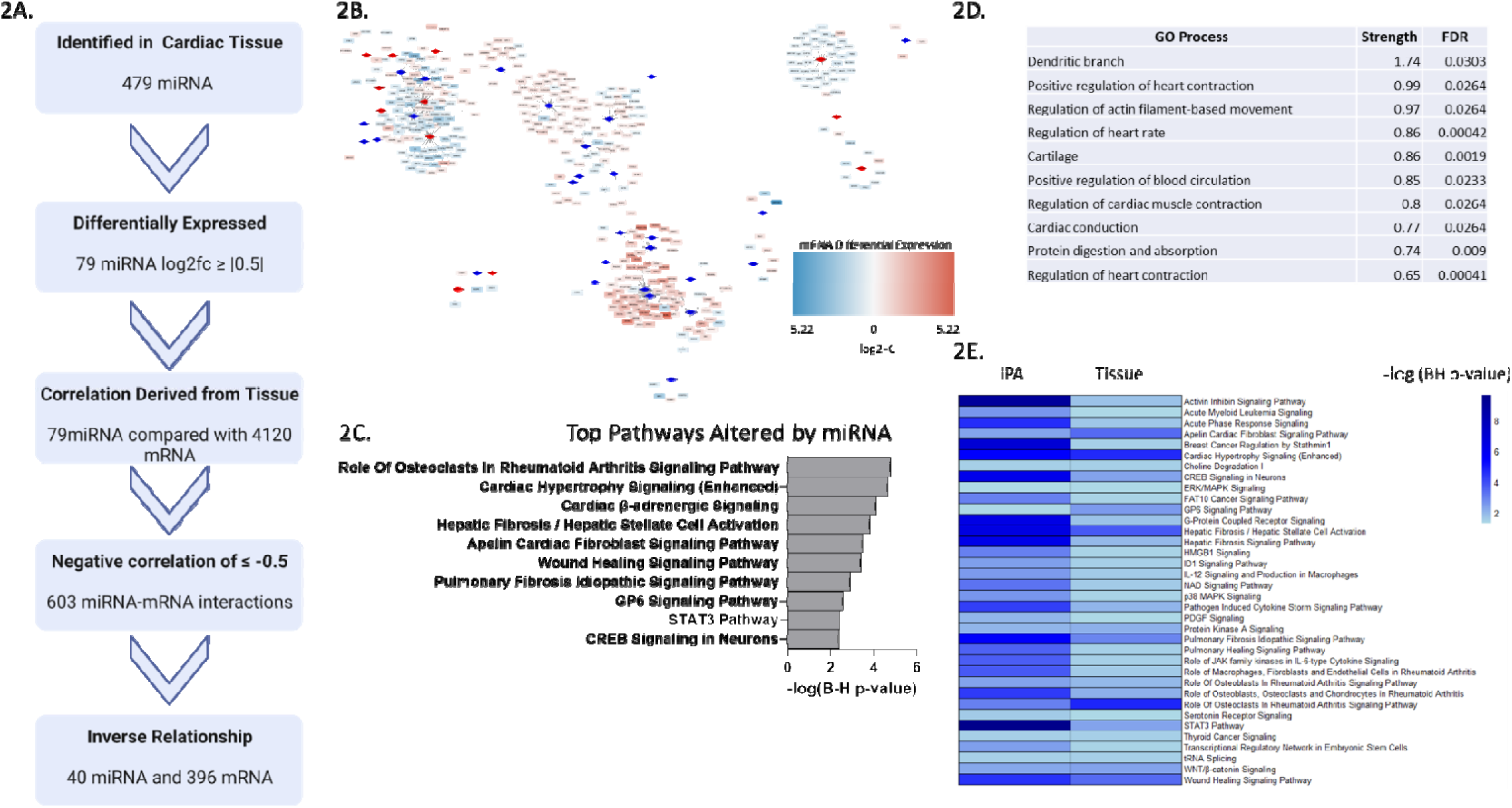
miRNA integration with transcriptome using correlation analysis. A) Integration workflow. B) mRNA-miRNA association network using correlation analysis. C) Top altered by miRNA. D) Enriched terms in mRNA-miRNA association network. E) Heatmap of pathways altered by miRNA utilizing both Ingenuity Pathway Analysis (IPA) and correlation analysis (Tissue).

Integrating the two methods showed 36 pathways which were confidently regulated by specific, differentially expressed miRNA including acute phase response signaling, cardiac hypertrophy signaling, ERK/MAPK signaling, hepatic fibrosis signaling, and p38 signaling (**Figure 2E**). Specifically, 8 miRNAs (miR-139-3p, miR-139-5p, miR-148a-3p, miR-92b-5p, miR-3065-5p, miR-184, miR-150-5p, and miR-144-5p) were associated with regulation of hypertrophy pathways using the correlation method (**Supplemental Figure 1B**). Thus, two methods, both an *in silico* database driven and an unbiased correlation analysis, suggest that a finite subset of miRNA regulate and may be ***sufficient*** to explain the transcriptional regulation of inflammatory, fibrotic, and hypertrophy pathways observed in HCM myectomy tissue.

Given that miRNA may explain the transcriptional downregulation of hypertrophy and inflammatory pathways, we next questioned whether miRNAs are ***necessary*** for these changes. To this end, an *in silico* knockout of the miRNA was performed. First, the miRNA-mRNA network generated using Ingenuity Pathway Analysis was used to identify DEGs impacted by miRNA. These DEGs were removed subsequently, and pathway analysis was performed on the remaining DEGs not impacted by miRNA to observe the pathways altered in HCM (**Figure 3A**). Following removal of the DEGs, of the 196 pathways altered in HCM samples, 79 (40%) pathways were no longer significantly altered, including numerous inflammatory and immune system related pathways (**Figure 3B and Supplemental Figure 2A**). Additionally, for 68 (35%) pathways the degree of dysregulation as measured by z-score was reduced. The hypertrophy pathways remained significantly altered however, the degree of downregulation was attenuated as well. For example, cardiac hypertrophy signaling (enhanced) (z-score -1.9 reduced to -1.0), ERK/MAPK (z-score -2.4 reduced to -1.5), and ERK-5 signaling (z-score -2.5 reduced to -2) were all still downregulated (**Figure 3C**). Interestingly, p38 MAPK, HER-2, IL-6, and STAT3 signaling were no longer altered.

**Figure 3.**
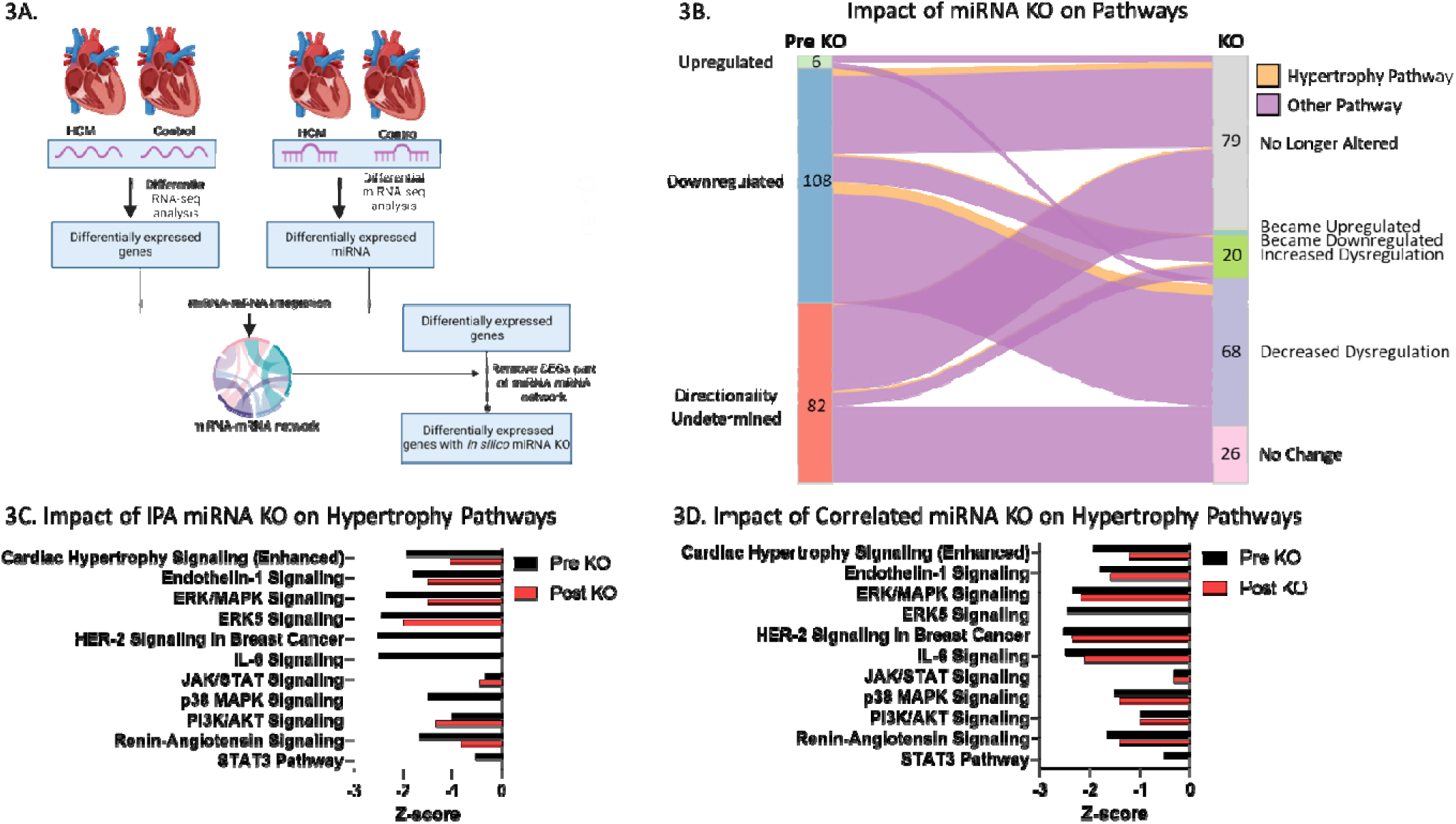
*In silico* knockout of miRNAs. A) Workflow to remove mRNA regulated by miRNA identified using Ingenuity Pathway Analysis. B) Alluvial plot showing changes in pathway status before (Pre KO) and after removing impact of miRNA (KO). C) Summary of changes in hypertrophy pathways before (Pre KO) and after removing impact of miRNA (KO) identified using Ingenuity Pathway Analysis. D) Summary of changes in hypertrophy pathways before (Pre KO) and after removing impact of miRNA (KO) identified using tissue correlation method. DEG, differentially expressed genes; KO, knock-out.

Again, to validate these *in silico* findings, the novel miRNA-mRNA regulatory network obtained from correlation analysis was used while the mRNA associated with miRNA was removed from the list of DEGs and pathway analysis was performed on the remaining DEG. Using this method 25/196 (13%) pathways were no longer significantly altered (**Supplemental Figure 2B**). There were 19 pathways that were knocked out by both methods. Similarly, the degree of downregulation of hypertrophy pathways was reduced but most of the hypertrophy pathways remained downregulated. For example, cardiac hypertrophy signaling (enhanced) (z-score -1.9 reduced to -1.2), ERK/MAPK (z-score -2.4 reduced to -2.1), and p38 MAPK signaling (z-score -1.5 reduced to -1.4) (**Figure 3D and Supplemental Figure 2B**). Interestingly, STAT3 and ERK5 signaling were no longer significantly altered using this method.

In summary, two independent miRNA-mRNA regulatory networks were generated to provide *in silico* evidence that the transcriptional dysregulation is partly attributed to a discrete set of miRNAs especially the downregulation of hypertrophy pathways and inflammatory pathways.

### Widespread dysregulation of histone architecture occurs in obstructive HCM including RAS-MAPK loci

To identify other possible modifiers of HCM transcriptional dysregulation, ChIP-seq was performed to identify differentially modified peaks for five different histone modifications (H3K27me3, H3K27ac, H3K9me3, H3K9ac, and H3K4me3) genome-wide (**Figure 4A**). **Supplemental Figures 3-7** provide a summary of the peak annotation for all five modifications. The list of differential peaks for each modification along with altered pathways are available in the **supplemental Excel file**. Only 853/31,734 (3%) H3K27me3 peaks were differentially altered between HCM and controls leading to significant enrichment of only 1 pathway: neuronal system (q-value = 0.04). Of the 138,293 H3K27ac peaks identified, 58,837 (43%) were differentially altered leading to enrichment of 50 pathways many of which were involved in RhoA and cytoskeletal signaling and MAPK signaling. Of the 64,060 H3K9me3 peaks identified only 33 (0.05%) were differentially modified. There were 23,339/109,921 (21%) H3K9ac peaks differentially modified impacting 87 pathways including Rho and MAPK pathways. Finally, across the 38,842 identified H3K4me3 peaks, 2912 (7%) were differentially methylated with 5 pathways impacted all involving Rho or Rac signaling. Thus, the HCM genome is characterized by widespread changes in histone modifications, whereby histone acetylation appears to be enriched in loci implicated in many pathways including RAS-MAPK signaling.

**Figure 4.**
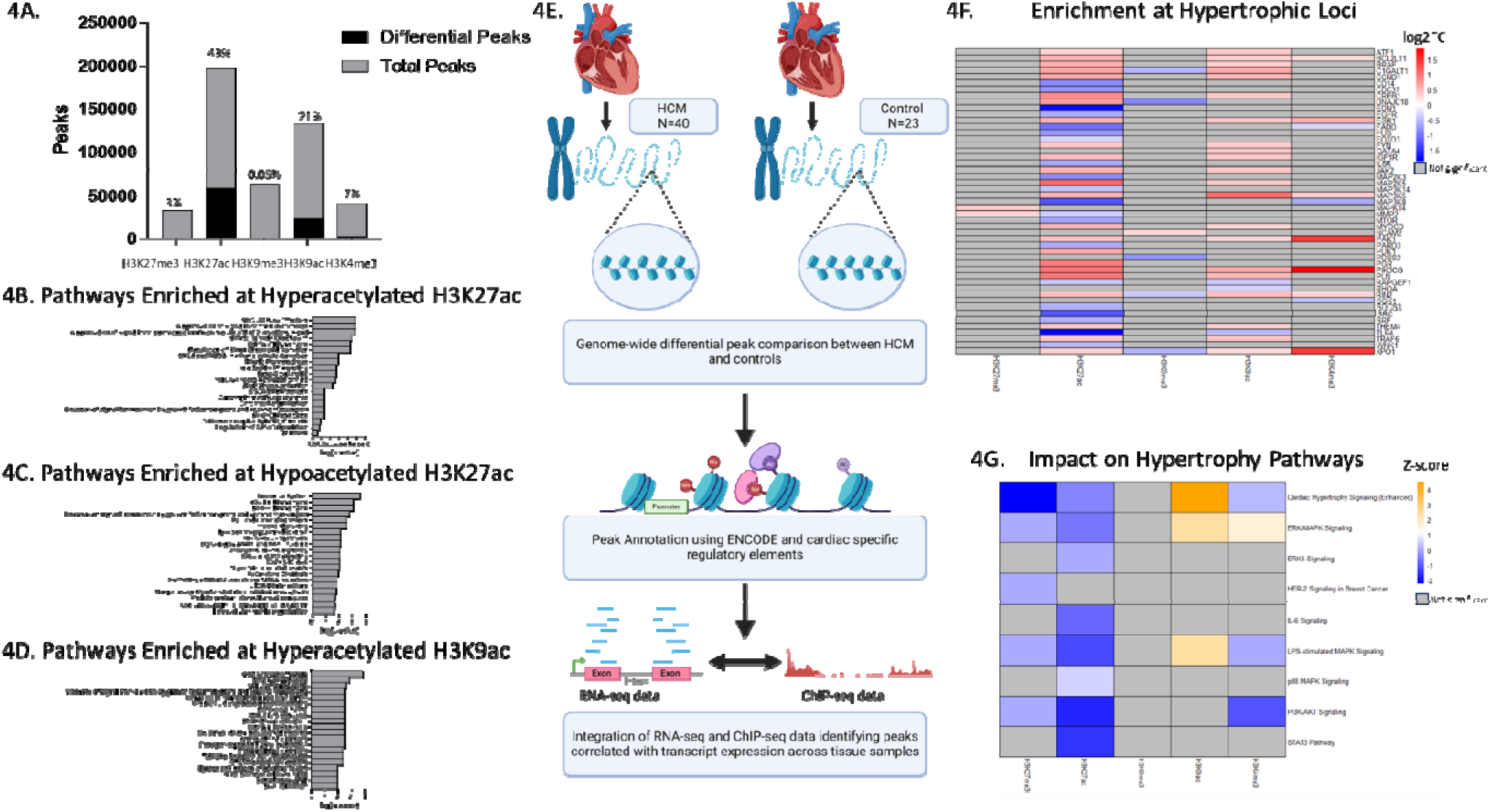
Comparison of post-translational modifications of histones between HCM and controls using chromatin immunoprecipitation sequencing (ChIP-seq). A) Summary of number of differential and total peaks identified for each modification. B) Pathways most enriched at hyperacetylated H3K27 loci. C) Pathways most enriched at hypoacetylated H3K27 loci. D) Pathways most enriched at hyperacetylated H3K9 loci. E) Workflow to integrate chromatin immunoprecipitation with transcriptome. F) Heatmap showing log2 fold change for hypertrophy-associated transcripts correlated with ChIP-seq loci. G) Summary of impact of ChIP-seq peaks correlated with transcripts on hypertrophy pathways. Not significant indicates loci/pathway was not enriched for that modification when integrating ChIP-seq with transcriptome. Ac, acetylation; log2fc, log2 fold change.

To test whether the enrichment of MAPK signaling loci was due to increased or decreased acetylation of H3K27 and H3K9, the increased and decreased loci for each modification were analyzed separately to identify enriched pathways (**Supplemental Excel file**). Performing pathway enrichment analysis on hyperacetylated H3K27 loci revealed 37 enriched pathways including MAP kinase activation (q-value = 0.008; **Figure 4B**) while 51 pathways were enriched in the hypoacetylated loci including signaling by BRAF and RAF1 fusions (q-value = 0.007) and oncogenic MAPK signaling (q-value = 0.008; **Figure 4C**). When looking only at H3K9 with increased acetylation, 94 pathways were enriched including MAP kinase activation (q-value = 0.003) and RAF activation (q-value = 0.01; **Figure 4D**) while none were enriched in the hypoacetylated loci. Thus, there is a global increase in acetylation of H3K9 loci at RAS-MAPK loci while there is a mixture of hypoacetylation and hyperacetylation of H3K27 at RAS-MAPK loci.

### Integration of histone modifications with the transcriptome reveals histone acetylation and methylation are associated with regulation of hypertrophy pathways

Next, the ChIP-seq data were integrated with the RNA-seq data to determine the impact of histone modifications on the transcriptome (**Figure 4E**). The differentially modified peaks were annotated using ENCODE and known distal cardiac regulatory elements to link differential peaks to specific transcripts. Correlation analysis was performed for each peak-transcript association and only ChIP-seq peaks statistically associated (adjusted p-value ≤ 0.05) with the corresponding transcripts’ expression in the tissue were considered if the directionality matched that expected for each modification (i.e., modifications associated with increased expression were directly correlated while those associated with decreased expression were inversely correlated; **Supplemental Excel file**). For each histone modification, pathway analysis was performed on the unique transcripts (**Supplemental Figure 8** and **Supplemental Excel file**). A total of 144 H3K27me3 peaks were inversely correlated with expression of 140 unique transcripts impacting 116 pathways. There were 9787 H3K27ac peaks directly correlated with expression of 2864 unique transcripts impacting 142 pathways. Only 7 H3K9me3 peaks were inversely correlated with expression of 7 unique transcripts impacting 5 pathways. While 4343 H3K9ac peaks were directly correlated with expression of 1695 unique transcripts impacting 120 pathways. Finally, 451 H3K4me3 peaks were directly correlated with expression of 368 unique transcripts impacting 68 pathways.

Interestingly, all 5 histone modifications were associated with expression of transcripts correlating to the hypertrophy pathways with H3K27ac correlated with the most hypertrophy transcripts (**Figure 4F**). Pathway analysis revealed H3K27me3 and K3K27ac were correlated with the most downregulation of hypertrophy pathways including cardiac hypertrophy signaling, ERK/MAPK signaling, ERK5 signaling, IL-6 signaling, LPS-stimulated MAPK signaling, and p38 MAPK signaling while H3K9ac was interestingly associated with upregulation of cardiac hypertrophy signaling and ERK/MAPK signaling (**Figure 4G**).

### Removing the impact of histone modifications on the transcriptome attenuates downregulation of hypertrophy pathways

Since histone modifications appear to regulate many pathways in HCM including several hypertrophy pathways, the next step was to abrogate the impact of the histone modifications to see if they are **necessary** for the transcriptional alterations observed. The transcripts regulated by histone modifications were removed from the list of DEGs and pathway analysis was performed (**Figure 5A**). Abolishing the effect of histone modifications changed 99/196 (50%) pathways to no longer be altered in the HCM transcriptome while 53 (27%) had reduced dysregulation (**Figure 5B and Supplemental Figure 9A**). Interestingly many of the hypertrophy pathways remained significantly altered but the degree of downregulation was reduced. For example, downregulation of cardiac hypertrophy signaling (enhanced) (z-score--1.9 to -1.7) and ERK/MAPK signaling (z-score--2.4 to -0.7) were both attenuated. Interestingly two hypertrophy pathways, ERK5 and p38 MPAK signaling were no longer altered when the effect of histones was removed.

**Figure 5.**
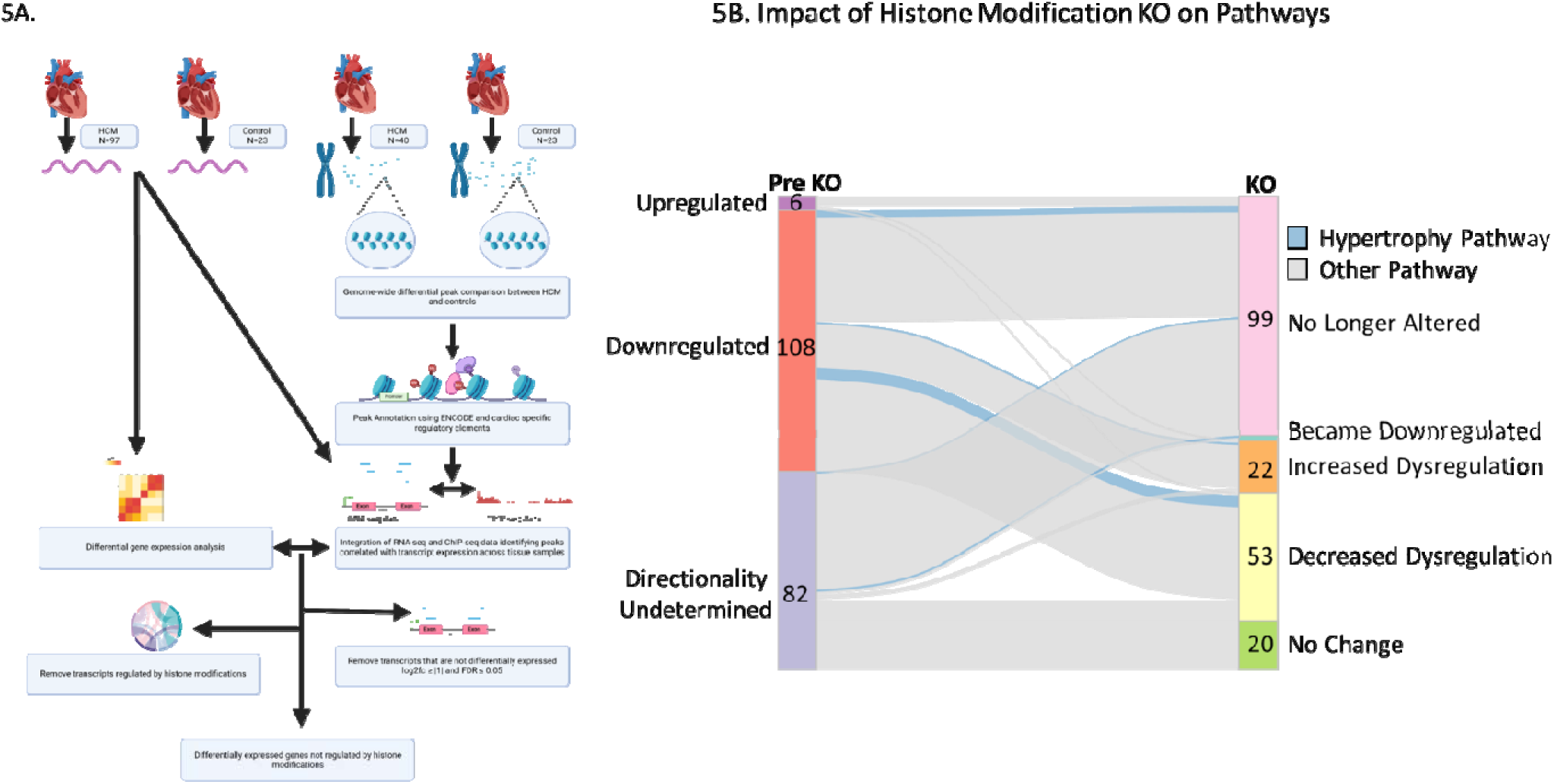
*In silico* knockout of histone modifications on the HCM transcriptome. A) Workflow to remove transcripts regulated by histone modifications. B) Alluvial plot showing changes in pathway status before (Pre KO) and after removing impact of histone modifications (KO).

### miRNA and Histone Modifications Act in Concert to Regulate Transcriptional Dysregulation in HCM

Finally, to see the impact of removing both the effect of miRNA and histones, all transcripts regulated by either the delimited set of differentially regulated miRNAs (using IPA method) and/or the histone modifications were removed from the list of DEGs and pathway analysis was performed resulting in 98/196 (50%) of pathways from being significantly altered to not being altered (**Figure 6A and Supplemental Figure 9B**). While removing both the effects of both miRNA and histone modifications did not drastically change the number of pathways impacted, it did abolish the transcriptional downregulation of cardiac hypertrophy signaling.

**Figure 6.**
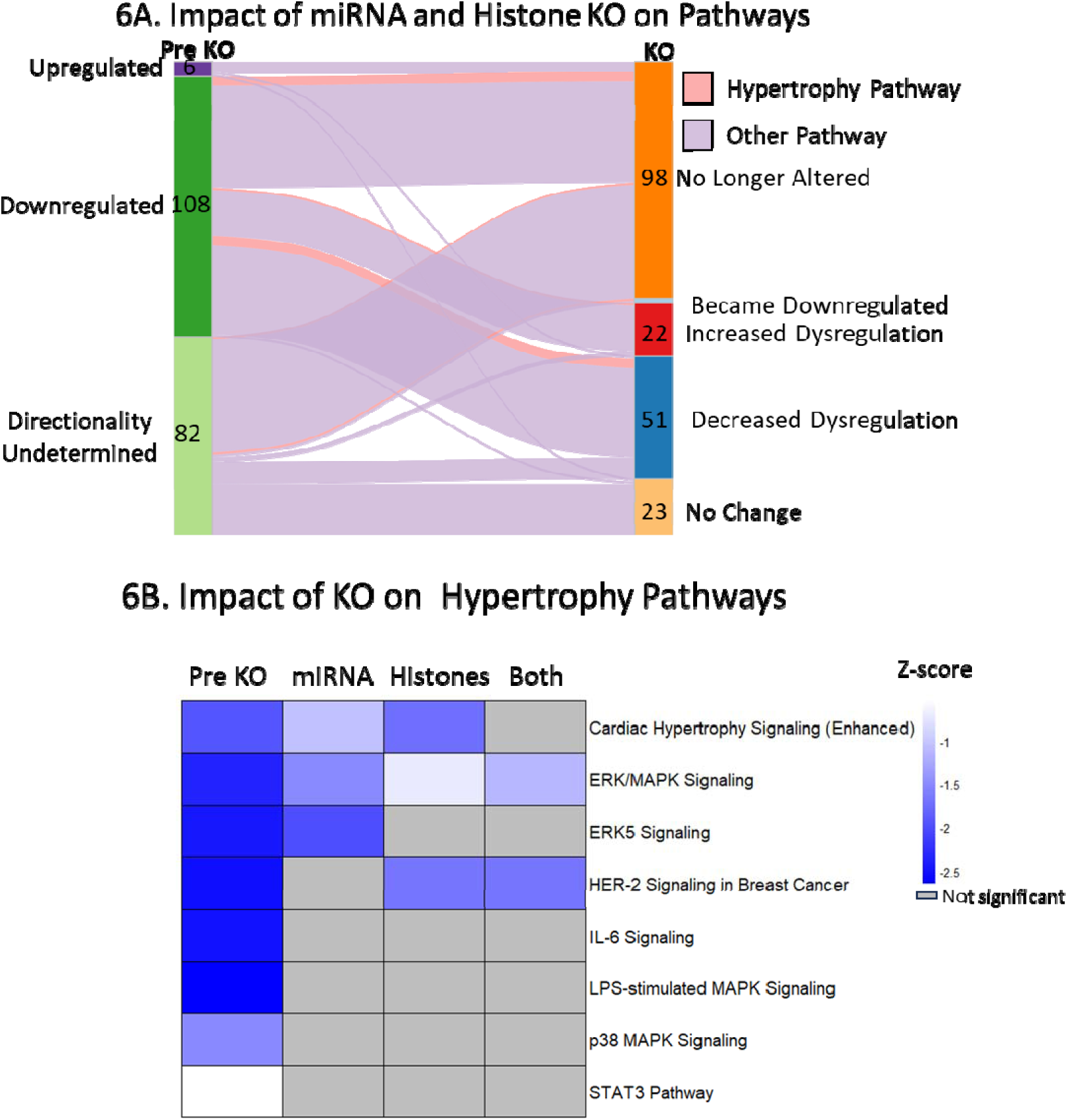
*In silico* knockout of miRNA and histone modifications on HCM transcriptome. A) Alluvial plot showing changes in pathway status before (Pre KO) and after removing impact of both miRNA and histone modifications (KO). B) Heatmap summarizing the impact of different *in silico* knockouts on status of hypertrophy pathways.

Figure 6B summarizes the impact of each scenario on the hypertrophy pathways. In summary, all hypertrophy pathways were downregulated transcriptionally when comparing HCM with controls. Abolishing the impact of either miRNA or histone modifications greatly reduced the transcriptional downregulation of hypertrophy pathways and removing the impact of both together almost completely removed the transcriptional downregulation of all hypertrophy pathways except ERK/MAPK signaling which was still downregulated but greatly reduced (z-score- -2.4 to -1.1). Thus late-stage obstructive HCM is characterized by transcriptional downregulation of hypertrophy pathways mediated in part by post-translational modifications in histones as well as alterations in miRNAs (Figure 7).

**Figure 7.**
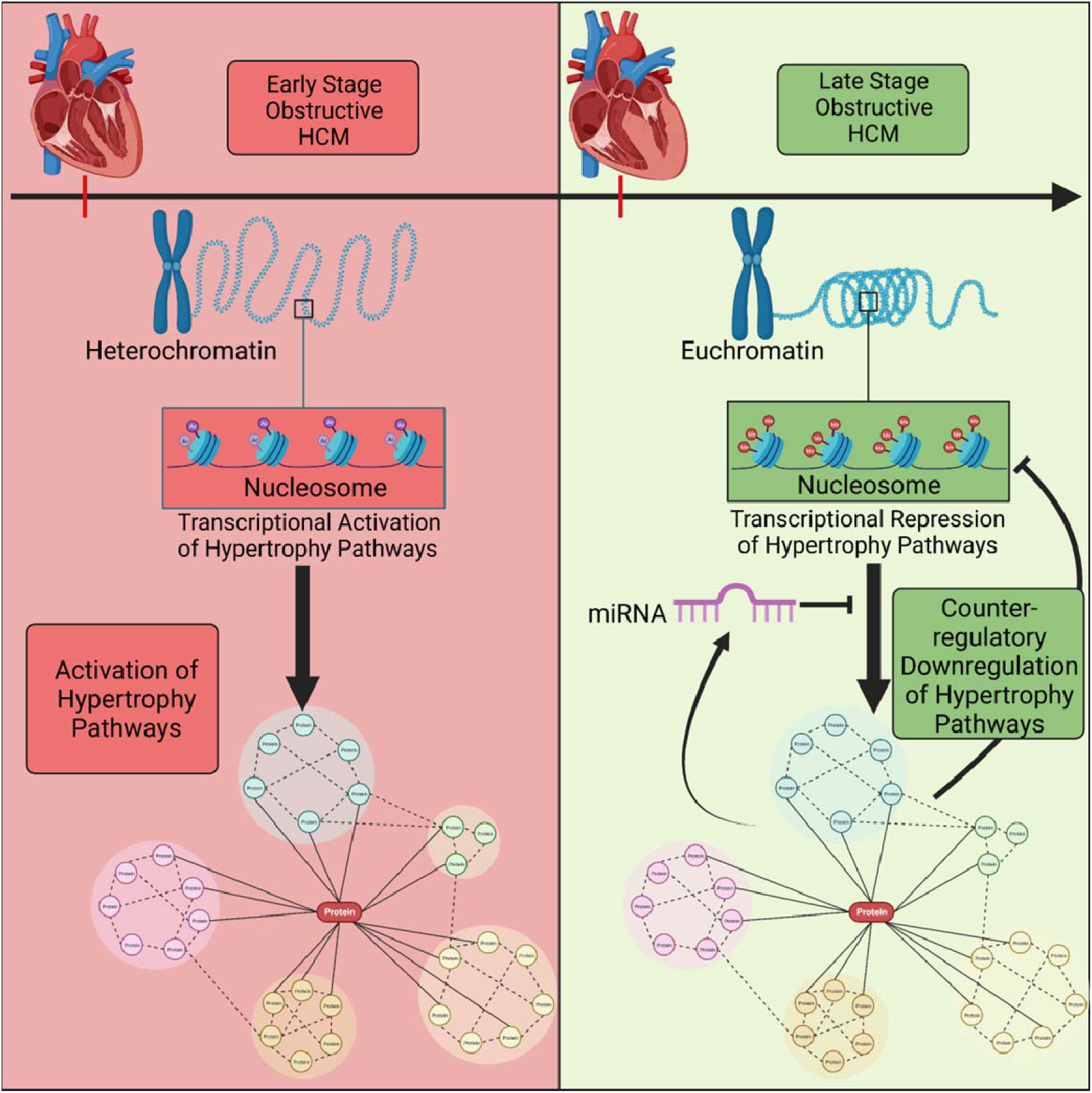
Proposed Model of Regulation of Hypertrophy Pathways in Obstructive HCM. In early disease we hypothesize there is transcriptional activation of hypertrophy pathways which leads to proteomic upregulation of hypertrophy pathways. In late-stage disease, we demonstrated continued translational upregulation of hypertrophy pathways with a counter-regulatory downregulation of hypertrophy pathways mediated by post-translational modifications in histones and miRNAs.

## Discussion

### A small subset of miRNAs are sufficient but not necessary to cause transcriptional dysregulation in HCM

To date, several studies have investigated differentially expressed miRNA in HCM plasma and myectomy samples^5–9, 28^. However, these studies used only pre-established interactions based on databases, and in most cases, there was no integration with RNA-sequencing data. Herein, we utilized two complementary approaches to develop two miRNA-mRNA regulatory networks by combining miRNA and mRNA sequencing data from the same HCM myectomy tissue. Both networks suggest that a small subset of miRNAs regulate fibrotic, inflammatory, and hypertrophy pathways in HCM. Specifically, we observed that miRNA regulated pathways previously implicated in cardiac hypertrophy and fibrosis, including RAS-MAPK, IL-6, PI3K/AKT, HER2, and STAT3 signaling ^29–37^.

Recently, Liang et al.^28^ similarly showed differentially expressed miRNA in plasma from HCM patients were predicted to regulate RAS-MAPK signaling. However, these finding were based on miRNA data alone. Our data showed that miRNA might indeed regulate the RAS-MAPK signaling, as identified in our previous study, and appear to be sufficient to result in transcriptional downregulation of hypertrophy pathways in obstructive HCM. While the miRNA by itself could result in downregulation of hypertrophy pathways and many of the transcriptional changes, we showed through computational abolishment of the miRNA, it was not necessary for the observed changes.

Although miRNAs did not seem to be entirely responsible for the transcriptional changes observed in HCM, the two novel miRNA-mRNA networks do increase our knowledge on pathogenesis HCM and could serve as a blueprint for development of therapeutic approaches.

For example, specific miRNA could be identified from the networks and potentially targeted and tested for their impact in altering the hypertrophy and fibrotic pathways early in disease.

### Role of histone modifications on the HCM transcriptome

Post-translational modifications of histones are known to play a crucial role in both normal and pathologic cardiac processes^38, 39^. Several studies have demonstrated that histone modifications (especially acetylation) play a role in other forms of cardiac hypertrophy such as pressure overload hypertrophy^12, 15, 39–41^. Despite this, except for only two small studies, the impact of histone modifications in HCM has been relatively unexplored^42, 43^. The first study correlated histone modifications to QRS associated loci while a second study looked at characterizing the role of histone modifications in *MYBPC3*-mediated HCM using a multi-omics approach in a small number of myectomy samples to demonstrate widespread changes in H3K27 acetylation and identify potential upstream regulators^43^.

Herein, we present our data on a much larger HCM cohort comprised of all the major HCM causative genotypes as well as genotype-negative individuals for 5 different histone modifications. Our histone marks integrated with our transcriptome data demonstrated for the first time in HCM that a large portion of the transcriptional dysregulation, including changes in cardiac hypertrophy and fibrotic pathways, may be due to changes in histone modifications.

Furthermore, using computational knock-out scenarios akin to our miRNA analyses described above, we show that removing the observed effect of histone marks abolishes much of the dysregulation, including the downregulation of fibrotic and hypertrophy pathways. Interestingly, we noticed a balance between hyper- and hypo-acetylation across hypertrophy-associated loci with H3K9ac appearing to result in transcriptional activation of hypertrophy pathways. H3K27ac correlated to both activation and repression of hypertrophy loci but overall leading to a repressive effect. Thus, in the myectomy tissue, there appears to be a balance between activation and repression of hypertrophy loci via histone modifications which in conjunction with miRNA leads to overall transcriptional downregulation of hypertrophy pathways.

### A role for epigenetic-targeted drugs as a novel future therapy for HCM?

One goal of this study was to illuminate the role of histone modifications in HCM and serve as a nidus for future studies looking at the functional role of epigenetics in HCM. The role of histone acetylation in cardiac hypertrophy is complex with a balance between regulation of pro- and anti-hypertrophy loci^13, 39^. Therapeutically, the use of histone deacetylase inhibitors (HDACis) can modulate the activity of histone deacetylases attenuating pathologic hypertrophy^13, 39^ and may therefore hold potential for the treatment of HCM. In fact, one study using an induced pluripotent stem cell-derived cardiomyocyte HCM model showed that HDACis reduced cellular hypertrophy and calcium dysregulation^44^. Interestingly, pressure overload hypertrophy was shown to activate target genes via increased H3K9ac and inducing deacetylation of target genes via HDACis attenuated the hypertrophy^13^. Myectomy tissue represents late-stage disease after significant hypertrophy has occurred^45^. We observed activation of hypertrophy pathways via increased H3K9ac but this was balanced by repression via miRNA, decreased H3K27ac, and increased H3K27me3.

Since histone modifications appear to play a role in transcriptional dysregulation in late-stage disease and may even have competing roles, future studies are necessary to identify the role of histone modifications in early HCM and the dynamic changes that occur as disease progresses. It would be important to see if early in disease, there are activation marks such as increased H3K9ac on hypertrophy loci similar to that seen with pressure overload hypertrophy leading to transcriptional activation of hypertrophy and disease pathways.

### Study Limitations and Future Directions

Studying HCM represents unique challenges with strong ethical and logistic barriers to acquiring cardiac tissue to study disease biology. Herein, we utilized one of the largest HCM myectomy tissue banks to characterize the epigenetic and miRNA architecture of HCM in myectomy tissue. While this provides a static view of clinically manifest, symptomatic obstructive HCM (later stage), it has provided critical insights in the pathogenesis and (in)active pathways of HCM. Future studies are necessary to identify the temporal progression of changes as well as to functionally validate the findings.

## Conclusion

Myectomy tissue from patients with obstructive HCM shows transcriptional dysregulation including transcriptional downregulation of hypertrophy pathways mediated by a small subset of miRNAs and post-translational modifications of histones. Cardiac hypertrophy loci showed activation via changes in H3K9ac and a mix of activation and repression via H3K27ac. Further studies are needed to understand the balance and temporal relationship between these competing regulators in HCM pathobiology and the potential for epigenetic therapies.

## Acknowledgements

We would like to thank the Mayo Clinic Genome Analysis Core and the Mayo Clinic Epigenomics Development Laboratory for their assistance in acquiring rigorous and high-quality data.

## Supplemental Materials

## References

1. Maron BJ and Maron MS. Hypertrophic cardiomyopathy. Lancet (London, England). 2013;381:242–55.

2. Garmany R, Bos JM, Tester DJ, Giudicessi JR, Dos Remedios C, Dasari S, Nagaraj NK, Nair AA, Johnson KL, Ryan ZC, Maleszewski JJ, Ommen SR, Dearani JA and Ackerman MJ. Multi-Omic Architecture of Obstructive Hypertrophic Cardiomyopathy. Circ Genom Precis Med. 2023.

3. Garmany R, Bos JM, Dasari S, Johnson KL, Tester DJ, Giudicessi JR, Dos Remedios C, Maleszewski JJ, Ommen SR, Dearani JA and Ackerman MJ. Proteomic and phosphoproteomic analyses of myectomy tissue reveals difference between sarcomeric and genotype-negative hypertrophic cardiomyopathy. Sci Rep. 2023;13:14341.

4. Lewis BP, Burge CB and Bartel DP. Conserved seed pairing, often flanked by adenosines, indicates that thousands of human genes are microRNA targets. Cell. 2005;120:15–20.

5. Sun D, Li C, Liu J, Wang Z, Liu Y, Luo C, Chen Y and Wen S. Expression Profile of microRNAs in Hypertrophic Cardiomyopathy and Effects of microRNA-20 in Inducing Cardiomyocyte Hypertrophy Through Regulating Gene MFN2. DNA and cell biology. 2019;38:796–807.

6. Wang L, Lu F and Xu J. Identification of Potential miRNA-mRNA Regulatory Network Contributing to Hypertrophic Cardiomyopathy (HCM). Frontiers in cardiovascular medicine. 2021;8:660372–660372.

7. Kuster DW, Mulders J, Ten Cate FJ, Michels M, Dos Remedios CG, da Costa Martins PA, van der Velden J and Oudejans CB. MicroRNA transcriptome profiling in cardiac tissue of hypertrophic cardiomyopathy patients with MYBPC3 mutations. Journal of molecular and cellular cardiology. 2013;65:59–66.

8. Li M, Chen X, Chen L, Chen K, Zhou J and Song J. MiR-1-3p that correlates with left ventricular function of HCM can serve as a potential target and differentiate HCM from DCM. Journal of translational medicine. 2018;16:161.

9. Osmak G, Baulina N, Kiselev I and Favorova O. MiRNA-Regulated Pathways for Hypertrophic Cardiomyopathy: Network-Based Approach to Insight into Pathogenesis. Genes. 2021;12.

10. Qin J, Guo N, Tong J and Wang Z. Function of histone methylation and acetylation modifiers in cardiac hypertrophy. Journal of molecular and cellular cardiology. 2021;159:120–129.

11. Mahmoud SA and Poizat C. Epigenetics and chromatin remodeling in adult cardiomyopathy. The Journal of pathology. 2013;231:147–57.

12. Liu CF and Tang WHW. Epigenetics in Cardiac Hypertrophy and Heart Failure. JACC Basic Transl Sci. 2019;4:976–993.

13. Ooi JY, Tuano NK, Rafehi H, Gao XM, Ziemann M, Du XJ and El-Osta A. HDAC inhibition attenuates cardiac hypertrophy by acetylation and deacetylation of target genes. Epigenetics. 2015;10:418–30.

14. Watson CJ, Horgan S, Neary R, Glezeva N, Tea I, Corrigan N, McDonald K, Ledwidge M and Baugh J. Epigenetic Therapy for the Treatment of Hypertension-Induced Cardiac Hypertrophy and Fibrosis. Journal of cardiovascular pharmacology and therapeutics. 2016;21:127–37.

15. Kook H, Lepore JJ, Gitler AD, Lu MM, Wing-Man Yung W, Mackay J, Zhou R, Ferrari V, Gruber P and Epstein JA. Cardiac hypertrophy and histone deacetylase-dependent transcriptional repression mediated by the atypical homeodomain protein Hop. J Clin Invest. 2003;112:863–71.

16. Papait R, Serio S and Condorelli G. Role of the Epigenome in Heart Failure. Physiological reviews. 2020;100:1753–1777.

17. Luger K, Mäder AW, Richmond RK, Sargent DF and Richmond TJ. Crystal structure of the nucleosome core particle at 2.8 A resolution. Nature. 1997;389:251–60.

18. Gray SG and Teh BT. Histone acetylation/deacetylation and cancer: an “open” and “shut” case? Current molecular medicine. 2001;1:401–29.

19. Black JC, Van Rechem C and Whetstine JR. Histone lysine methylation dynamics: establishment, regulation, and biological impact. Molecular cell. 2012;48:491–507.

20. Caride A, Jang JS, Shi GX, Lenz S, Zhong J, Kim KH, Allen M, Robertson KD, Farrugia G, Ordog T, Ertekin-Taner N and Lee JH. Titration-based normalization of antibody amount improves consistency of ChIP-seq experiments. BMC Genomics. 2023;24:171.

21. Kechin A, Boyarskikh U, Kel A and Filipenko M. cutPrimers: A New Tool for Accurate Cutting of Primers from Reads of Targeted Next Generation Sequencing. Journal of computational biology : a journal of computational molecular cell biology. 2017;24:1138–1143.

22. Li H and Durbin R. Fast and accurate short read alignment with Burrows-Wheeler transform. Bioinformatics. 2009;25:1754–60.

23. Zhang Y, Liu T, Meyer CA, Eeckhoute J, Johnson DS, Bernstein BE, Nusbaum C, Myers RM, Brown M, Li W and Liu XS. Model-based analysis of ChIP-Seq (MACS). Genome biology. 2008;9:R137.

24. Zang C, Schones DE, Zeng C, Cui K, Zhao K and Peng W. A clustering approach for identification of enriched domains from histone modification ChIP-Seq data. Bioinformatics. 2009;25:1952–8.

25. Amemiya HM, Kundaje A and Boyle AP. The ENCODE Blacklist: Identification of Problematic Regions of the Genome. Sci Rep. 2019;9:9354.

26. Ross-Innes CS, Stark R, Teschendorff AE, Holmes KA, Ali HR, Dunning MJ, Brown GD, Gojis O, Ellis IO, Green AR, Ali S, Chin SF, Palmieri C, Caldas C and Carroll JS. Differential oestrogen receptor binding is associated with clinical outcome in breast cancer. Nature. 2012;481:389–93.

27. Yu G, Wang LG and He QY. ChIPseeker: an R/Bioconductor package for ChIP peak annotation, comparison and visualization. Bioinformatics. 2015;31:2382–3.

28. Liang LW, Hasegawa K, Maurer MS, Reilly MP, Fifer MA and Shimada YJ. Comprehensive Transcriptomics Profiling of MicroRNA Reveals Plasma Circulating Biomarkers of Hypertrophic Cardiomyopathy and Dysregulated Signaling Pathways. Circ Heart Fail. 2023;16:e010010.

29. Page C and Doubell AF. Mitogen-Activated Protein Kinase (MAPK) in Cardiac Tissues. Mol Cell Biochem. 1996;157:49–57.

30. Robinson P, Liu X, Sparrow A, Patel S, Zhang YH, Casadei B, Watkins H and Redwood C. Hypertrophic Cardiomyopathy Mutations increase Myofilament Ca(2+) Buffering, Alter Intracellular Ca(2+) Handling, and Stimulate Ca(2+)-Dependent Signaling. J Biol Chem. 2018;293:10487–10499.

31. Lombardi R, Rodriguez G, Chen SN, Ripplinger CM, Li W, Chen J, Willerson JT, Betocchi S, Wickline SA, Efimov IR and Marian AJ. Resolution of Established Cardiac Hypertrophy and Fibrosis and Prevention of Systolic Dysfunction in a Transgenic Rabbit Model of Human Cardiomyopathy Through Thiol-Sensitive Mechanisms. Circulation. 2009;119:1398–407.

32. Chou CH, Hung CS, Liao CW, Wei LH, Chen CW, Shun CT, Wen WF, Wan CH, Wu XM, Chang YY, Wu VC, Wu KD and Lin YH. IL-6 Trans-signalling Contributes to Aldosterone-Induced Cardiac Fibrosis. Cardiovasc Res. 2018;114:690–702.

33. Nicol RL, Frey N, Pearson G, Cobb M, Richardson J and Olson EN. Activated MEK5 Induces Serial Assembly of Sarcomeres and Eccentric Cardiac Hypertrophy. Embo j. 2001;20:2757–67.

34. Razzaque MA, Nishizawa T, Komoike Y, Yagi H, Furutani M, Amo R, Kamisago M, Momma K, Katayama H, Nakagawa M, Fujiwara Y, Matsushima M, Mizuno K, Tokuyama M, Hirota H, Muneuchi J, Higashinakagawa T and Matsuoka R. Germline Gain-of-Function Mutations in RAF1 Cause Noonan Syndrome. Nat Genet. 2007;39:1013–7.

35. Shiojima I, Yefremashvili M, Luo Z, Kureishi Y, Takahashi A, Tao J, Rosenzweig A, Kahn CR, Abel ED and Walsh K. Akt Signaling Mediates Postnatal Heart Growth in Response to insulin and Nutritional Status. J Biol Chem. 2002;277:37670–7.

36. Chan HW, Jenkins A, Pipolo L, Hannan RD, Thomas WG and Smith NJ. Effect of dominant-negative epidermal growth factor receptors on cardiomyocyte hypertrophy. Journal of receptor and signal transduction research. 2006;26:659–77.

37. Ye S, Luo W, Khan ZA, Wu G, Xuan L, Shan P, Lin K, Chen T, Wang J, Hu X, Wang S, Huang W and Liang G. Celastrol Attenuates Angiotensin II-Induced Cardiac Remodeling by Targeting STAT3. Circ Res. 2020;126:1007–1023.

38. Zhao Y and Garcia BA. Comprehensive Catalog of Currently Documented Histone Modifications. Cold Spring Harbor perspectives in biology. 2015;7:a025064.

39. Backs J and Olson EN. Control of cardiac growth by histone acetylation/deacetylation. Circ Res. 2006;98:15–24.

40. Burridge PW, Sharma A and Wu JC. Genetic and Epigenetic Regulation of Human Cardiac Reprogramming and Differentiation in Regenerative Medicine. Annual review of genetics. 2015;49:461–84.

41. Papait R, Cattaneo P, Kunderfranco P, Greco C, Carullo P, Guffanti A, Viganò V, Stirparo GG, Latronico MV, Hasenfuss G, Chen J and Condorelli G. Genome-wide analysis of histone marks identifying an epigenetic signature of promoters and enhancers underlying cardiac hypertrophy. Proc Natl Acad Sci U S A. 2013;110:20164–9.

42. Hemerich D, Pei J, Harakalova M, van Setten J, Boymans S, Boukens BJ, Efimov IR, Michels M, van der Velden J, Vink A, Cheng C, van der Harst P, Moore JH, Mokry M, Tragante V and Asselbergs FW. Integrative Functional Annotation of 52 Genetic Loci Influencing Myocardial Mass Identifies Candidate Regulatory Variants and Target Genes. Circ Genom Precis Med. 2019;12:e002328.

43. Pei J, Schuldt M, Nagyova E, Gu Z, El Bouhaddani S, Yiangou L, Jansen M, Calis JJA, Dorsch LM, Blok CS, van den Dungen NAM, Lansu N, Boukens BJ, Efimov IR, Michels M, Verhaar MC, de Weger R, Vink A, van Steenbeek FG, Baas AF, Davis RP, Uh HW, Kuster DWD, Cheng C, Mokry M, van der Velden J, Asselbergs FW and Harakalova M. Multi-omics integration identifies key upstream regulators of pathomechanisms in hypertrophic cardiomyopathy due to truncating MYBPC3 mutations. Clinical epigenetics. 2021;13:61.

44. Han L, Li Y, Tchao J, Kaplan AD, Lin B, Li Y, Mich-Basso J, Lis A, Hassan N, London B, Bett GC, Tobita K, Rasmusson RL and Yang L. Study familial hypertrophic cardiomyopathy using patient-specific induced pluripotent stem cells. Cardiovasc Res. 2014;104:258–69.

45. Ommen SR, Maron BJ, Olivotto I, Maron MS, Cecchi F, Betocchi S, Gersh BJ, Ackerman MJ, McCully RB, Dearani JA, Schaff HV, Danielson GK, Tajik AJ and Nishimura RA. Long-term effects of surgical septal myectomy on survival in patients with obstructive hypertrophic cardiomyopathy. Journal of the American College of Cardiology. 2005;46:470–6.

