## Supplemental Figures for "Histone Modifications and miRNA Perturbations Contribute to Transcriptional Dysregulation of Hypertrophy in Obstructive Hypertrophic Cardiomyopathy"

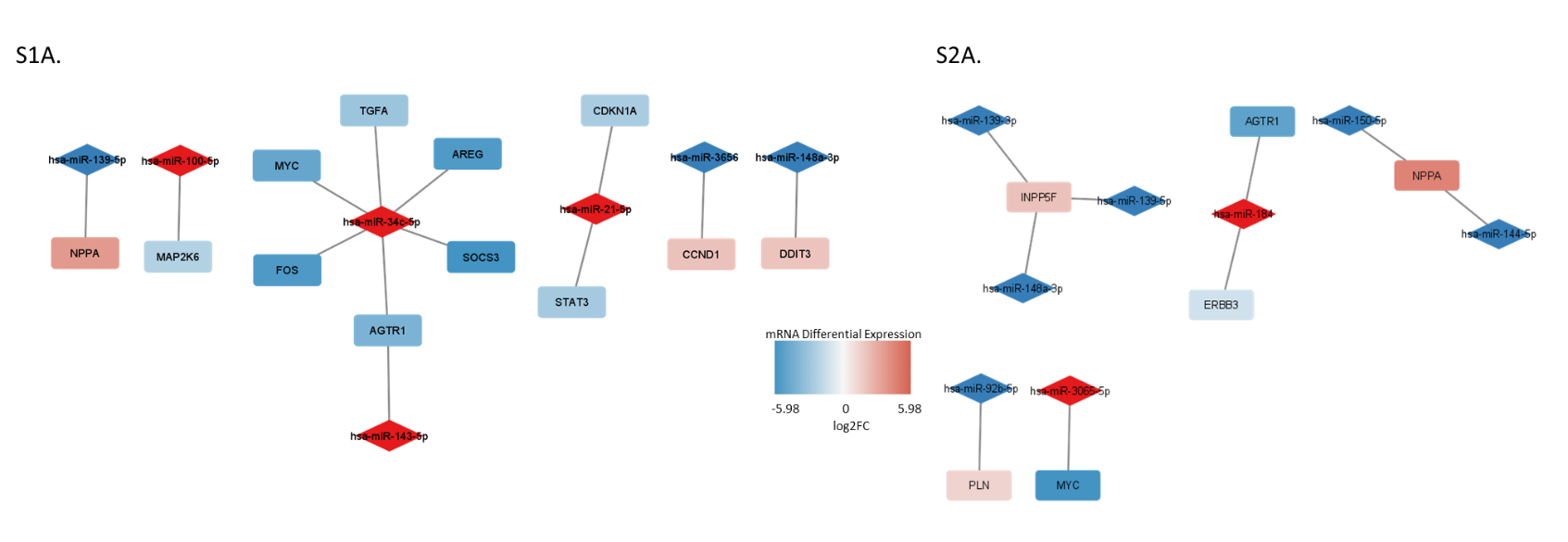

**Supplemental Figure 1.** Hypertrophy transcripts associated with miRNA A) using both Ingenuity Pathway Analysis method and B) using tissue correlation analysis.

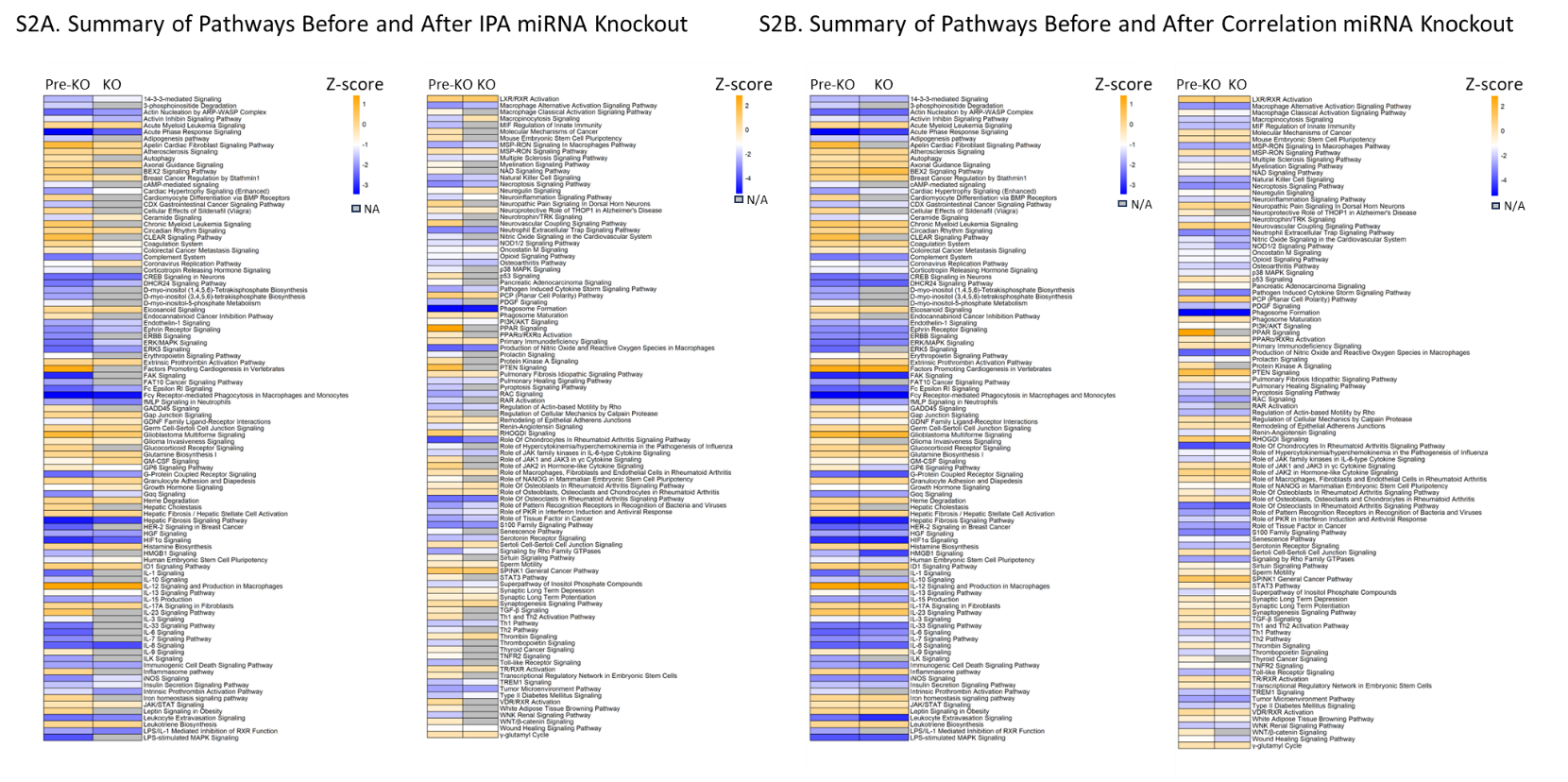
**Supplemental Figure 2.** Summary of pathway status before (Pre-KO) and after removing impact of miRNA (KO) using both Ingenuity Pathway Analysis (IPA) derived (A) and tissue correlation derived (B) miRNA. N/A indicates pathway is no longer altered when removing impact of miRNA. KO, knock-out.

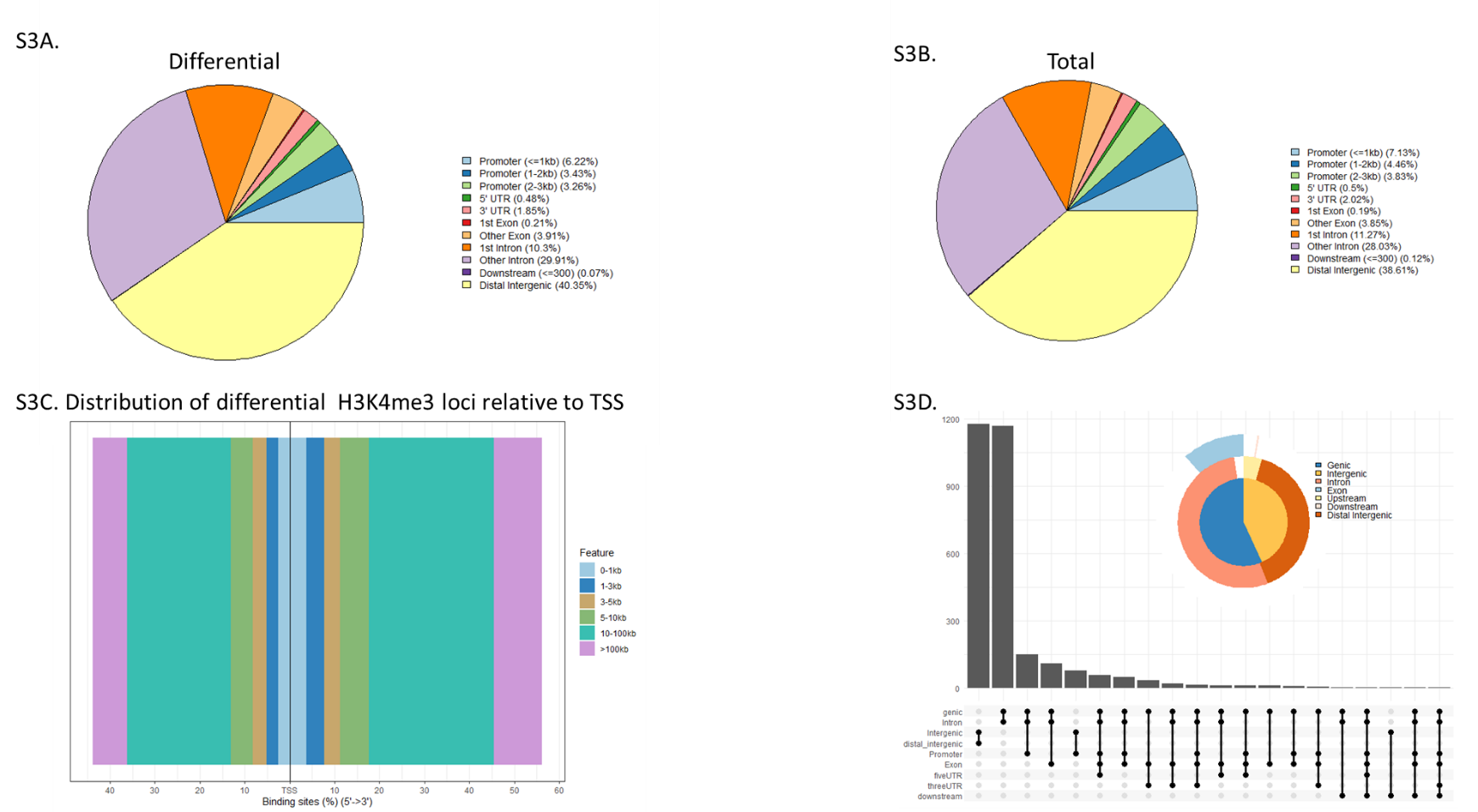

**Supplemental Figure 3.** H3K4me3 peak annotation using ChIPseeker. A) Circle plot of peak distribution for differential peaks. B) Circle plot of peak distribution for total peaks. C) Distribution plot of differential peaks relative to transcriptional start site (TSS). D) UpSet plot summarizing differential peak data.

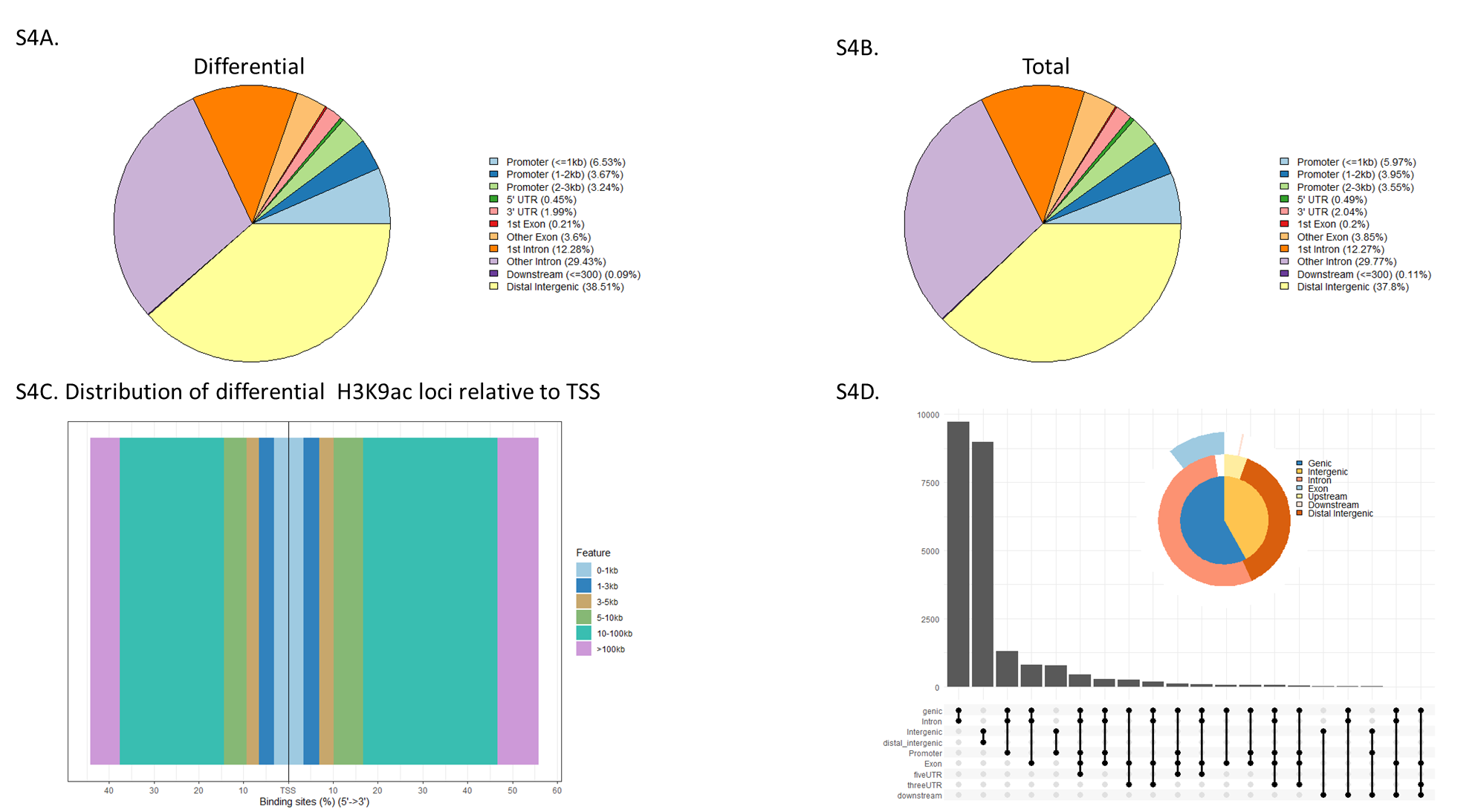

**Supplemental Figure 4.** H3K9ac peak annotation using ChIPseeker. A) Circle plot of peak distribution for differential peaks. B) Circle plot of peak distribution for total peaks. C) Distribution plot of differential peaks relative to transcriptional start site (TSS). D) UpSet plot summarizing differential peak data.

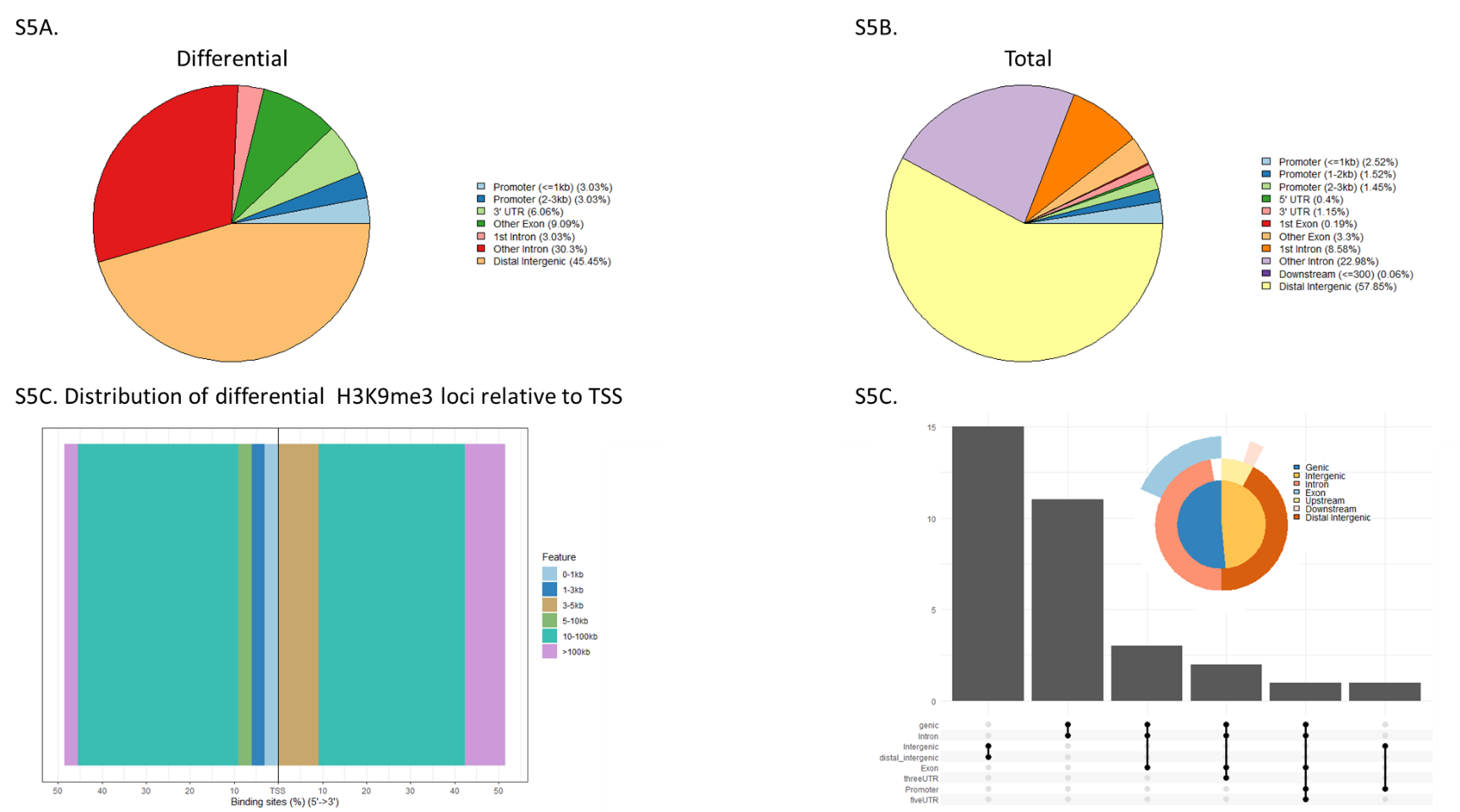

**Supplemental Figure 5.** H3K9me3 peak annotation using ChIPseeker. A) Circle plot of peak distribution for differential peaks. B) Circle plot of peak distribution for total peaks. C) Distribution plot of differential peaks relative to transcriptional start site (TSS). D) UpSet plot summarizing differential peak data.

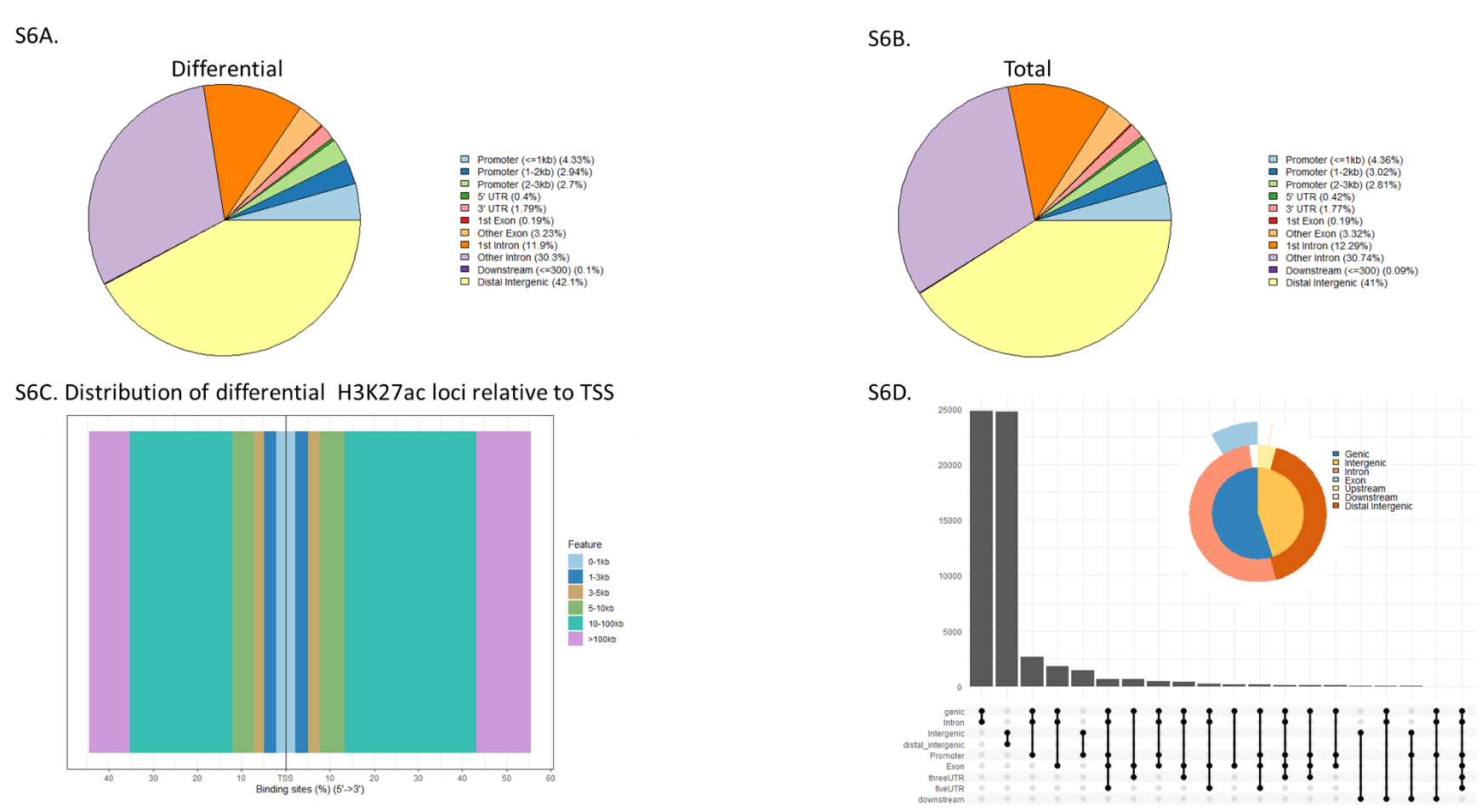

**Supplemental Figure 6.** H3K27ac peak annotation using ChIPseeker. A) Circle plot of peak distribution for differential peaks. B) Circle plot of peak distribution for total peaks. C) Distribution plot of differential peaks relative to transcriptional start site (TSS). D) UpSet plot summarizing differential peak data.

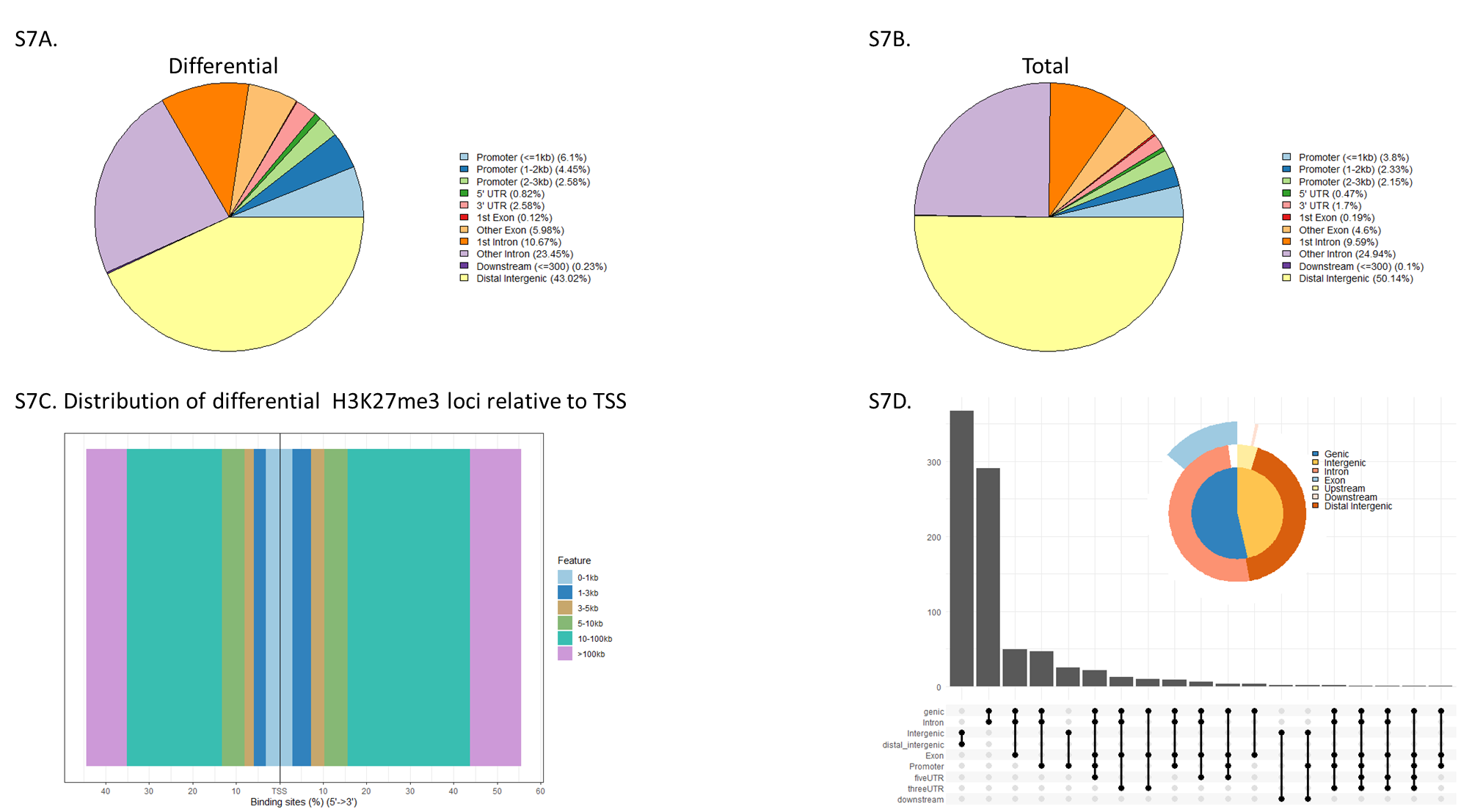

**Supplemental Figure 7.** H3K27me3 peak annotation using ChIPseeker. A) Circle plot of peak distribution for differential peaks. B) Circle plot of peak distribution for total peaks. C) Distribution plot of differential peaks relative to transcriptional start site (TSS). D) UpSet plot summarizing differential peak data.

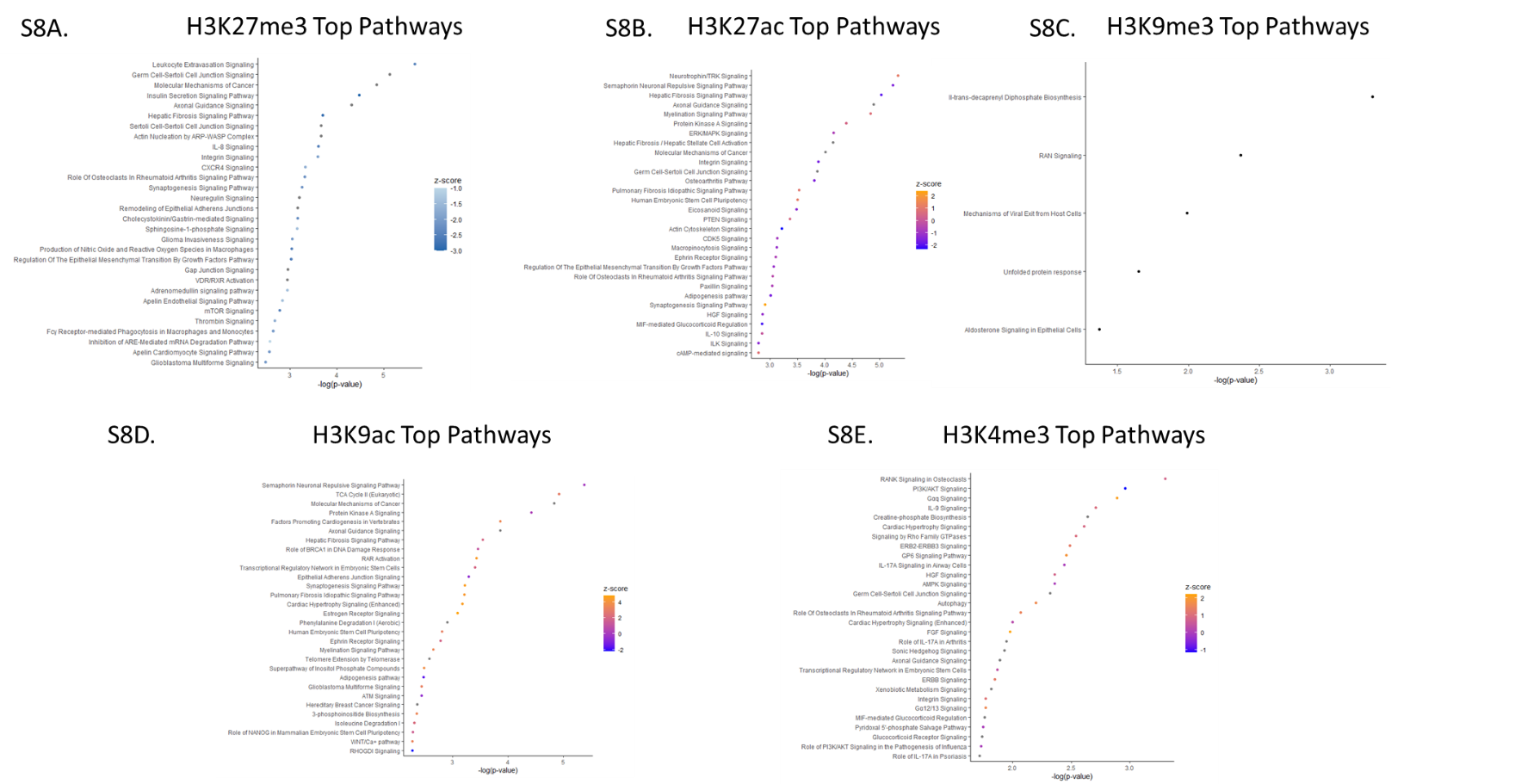

**Supplemental Figure 8.** Pathway enrichment of differential ChIP-seq peaks correlated with transcriptome. A) Top pathways enriched for H3K27me3. B) Top pathways enriched for H3K27ac. C) Top pathways enriched for H3K9me3. D) Top pathways enriched for H3K9ac. E) Top pathways enriched for H3K4me3.

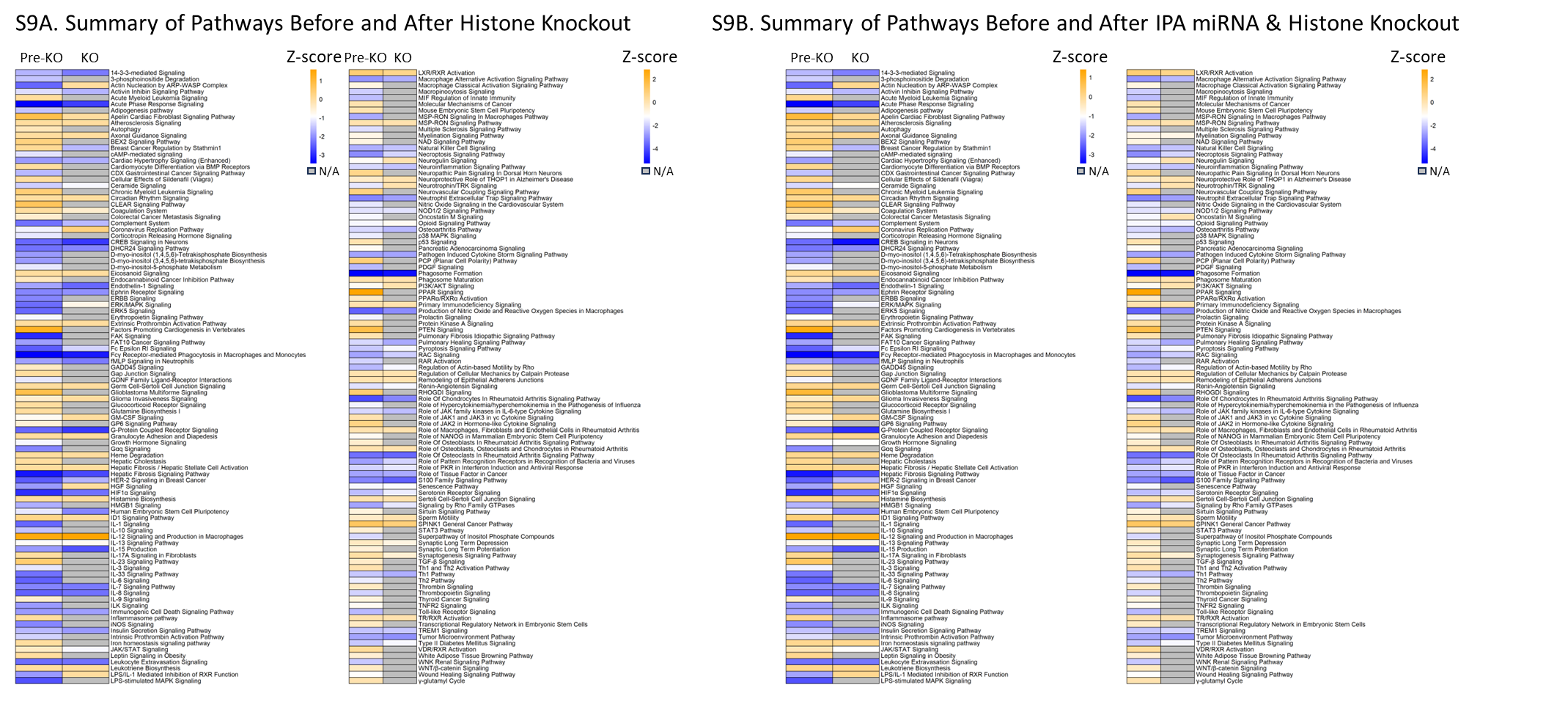

**Supplemental Figure 9.** Summary of pathway status before (Pre-KO) and after removing impact of histone modifications (KO) (A) and removing impact of both miRNA and histone modifications (B) miRNA. N/A indicates pathway is no longer altered when removing impact of miRNA. KO, knock-out.
